# Invasive Siberian chipmunks *Eutamias sibiricus* in Italy: a socio-ecological analysis indicates that they could, and should, be removed

**DOI:** 10.1101/535088

**Authors:** Jacopo Cerri, Emiliano Mori, Rudy Zozzoli, Andrea Gigliotti, Antony Chirco, Sandro Bertolino

## Abstract

**BACKGROUND:** Eradication of invasive alien species is a form of pest control linked to biodiversity conservation that usually involves killing animals. Squirrels are prominent among invasive alien species in Italy with four species introduced. Three of them are included within the list of alien species of European concern and their eradication and control is recommended. However, their local control is not an easy task, being highly appreciated by the general public. We propose a socio-ecological approach to evaluate the feasibility of eradicating Siberian chipmunks (*Eutamias sibiricus*) populations. We performed a structured questionnaire to assess the social perception of invasive Siberian chipmunks in urban parks where they occur and to identify groups of visitors who might oppose eradication. We also carried out geographic profiling to test for the spatial expansion of chipmunk populations.

**RESULTS:** Overall, park visitors regarded chipmunks positively and appreciated to see them, but human-chipmunk interactions were still rare. We did not identify any group of visitors with a strong attachment to chipmunks, who might oppose future control programs. Geographic profiling showed that chipmunks in Valeggio sul Mincio are starting to expand outside of their introduction site.

**CONCLUSIONS:** Data from questionnaires show that chipmunks eradication, coupled with adequate communication initiatives, might be feasible. Moreover, geographic profiling indicates that time for a rapid removal is running out. Socio-ecological approaches, combining the analysis of structured questionnaires administered to stakeholders and statistical modeling of pest observations, could be a valuable tool to decide the feasibility and the urgency of invasive pest control.

## 1 INTRODUCTION

The Siberian chipmunk *Eutamias sibiricus* is a widespread species in Russia and the Far East, which has become invasive in some European countries since the 1960s after its widespread trade as a pet species.^1^ Chipmunks established viable populations In Italy^1^, chipmunks that escaped from captivity established four viable populations in Northern Italy, and at two urban parks in Rome. Siberian chipmunks are not a mainstream invader, as the Eastern grey squirrel *Sciurus carolinensis*;^2^ however, they are considered an invasive alien species of European concern being listed within the EU Regulation 1143/2014. Particularly, chipmunks can act as a vector of tick-borne diseases and zoonoses.^3,4^ The European regulation requires member states to eradicated listed species from their territories when it is still possible and this appear the situation in Italy where the species is still localized with small populations.^5,6^

Management interventions aimed at containing or removing invasive alien mammals are more feasible when two ideal conditions occur. First, they are more cost-effective and face a higher success rate whenever the target species are still in the early stages of their invasion.^7-9^ Second, eradication initiatives tend to be more feasible when target species have minor interactions with society.^10^ Attempts to remove iconic alien mammals could result into strong opposition from some stakeholders.^11,12^ For instance, a trial eradication of the gray squirrel from Italy attempted in 1990s was stopped by animal right groups who brought the case in front of the court.^13^

Preliminary social impact assessments can tell managers whether control interventions will be opposed by relevant stakeholders and how they can be designed accordingly, to be successful. ^14,15^ Socio-ecological assessments go one step further, by combining information from relevant stakeholders, obtained through qualitative or quantitative methods from the social sciences, with information about the ecology, distribution and population dynamics of target species^16^. Ecological information can also be spatially explicit, as most ecological processes incorporate a geographical dimension^17^.

Eradicating chipmunks from Italy is required under the European and national legal framework, but, to the best of our knowledge, no study was conducted to verify if the preconditions for a successful eradication occur. Social prerequisites are particular sensitive considering the strong opposition faced by managers aiming to control or eradicate the gray squirrel in this country, even in recent years.^13, 18, 19^

Our research aims to fill this gap, by conduction a socio-ecological analysis combining spatial data of the species altogether with information from a structured surveys administered to a sample of visitors. Our analysis aims to test whether chipmunks became an iconic species and to inspect patterns in their geographical spread over time, to identify at which stage their invasion might be.

## 2 MATERIALS AND METHODS

In this research we focused on the three urban parks where chipmunks established viable populations in Italy. The first one is Sigurtà Garden Park, in Valeggio Sul Mincio, where Korean chipmunks were released in 1978, establishing the largest Italian populations of chipmunks in Italy.^1, 20-22^ The latter two areas where two urban parks in Rome, Villa Ada, where chipmunks were introduced at multiple times since the early 1908s and Villa Doria-Pamphilii, where chipmunks were observed for the first time in 2018.^6^

We surveyed a sample of visitors, administering a structured questionnaire measuring some their interactions with chipmunks and some psychological drivers of human-chipmunk interactions: attitudes, emotional dispositions, core affect, existence beliefs, social norms and behavioral intention about the presence of chipmunks. Attitudes were measured by means of a Likert scale and they were conceptualized as divided in some beliefs, characterized each one by its strength and the evaluation of its outcome.^23^ The attitudinal scale was built up by considering all the potential impacts of a species of ground squirrels living in a park, after a pilot study (S1). Emotions were measured as emotional disposition (joy, fear, surprise, disgust, interest) and core affect, or the extent respondents would have felt positive or negative at the idea of encountering a chipmunk.^24-27^ We measured existence beliefs by asking respondents to rate the importance of chipmunks in the park, both for future generations and per se.^28^ We measured social norms about the appropriateness of chipmunk presence in the park, by using three items measuring moral beliefs, empirical and normative expectations, and the willingness to enforce them by reporting the presence of chipmunks to local authorities.^29^ Visitors were also asked whether they had ever heard of chipmunks living in the park and if they had ever seen, fed or touched them. A complete list of the various questions adopted in the questionnaire, altogether with their summary is available in Table 1 and a complete copy of the questionnaire at (https://docs.google.com/forms/d/e/1FAIpQLSelkWacunZsTWDB3Qxy3QCYvfaFKg-hhd9LFmhcLcEZAbyfuA/viewform?usp=sf_link). The questionnaires were administered in Rome in both parks, but the collected data were grouped together as the two parks share the same pool of visitors. The questionnaire was implemented on GoogleForms. Most respondents (93.95%) were recruited on the field and they completed the questionnaire on a tablet. An online version was also administered on some Facebook groups on these urban parks. Questionnaires were confidential and they took approximately 15 minutes to be filled.

**Table 1.**
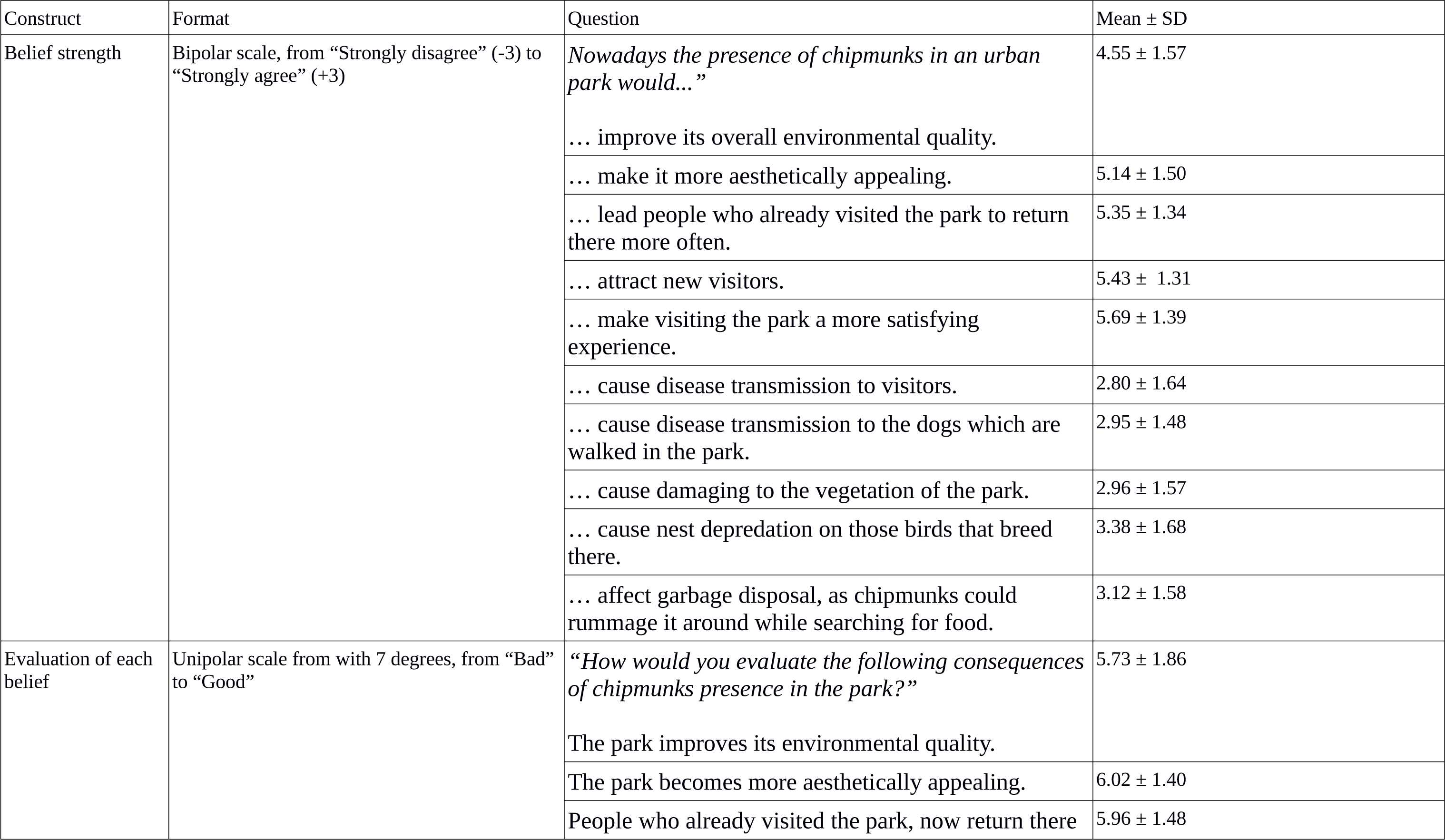
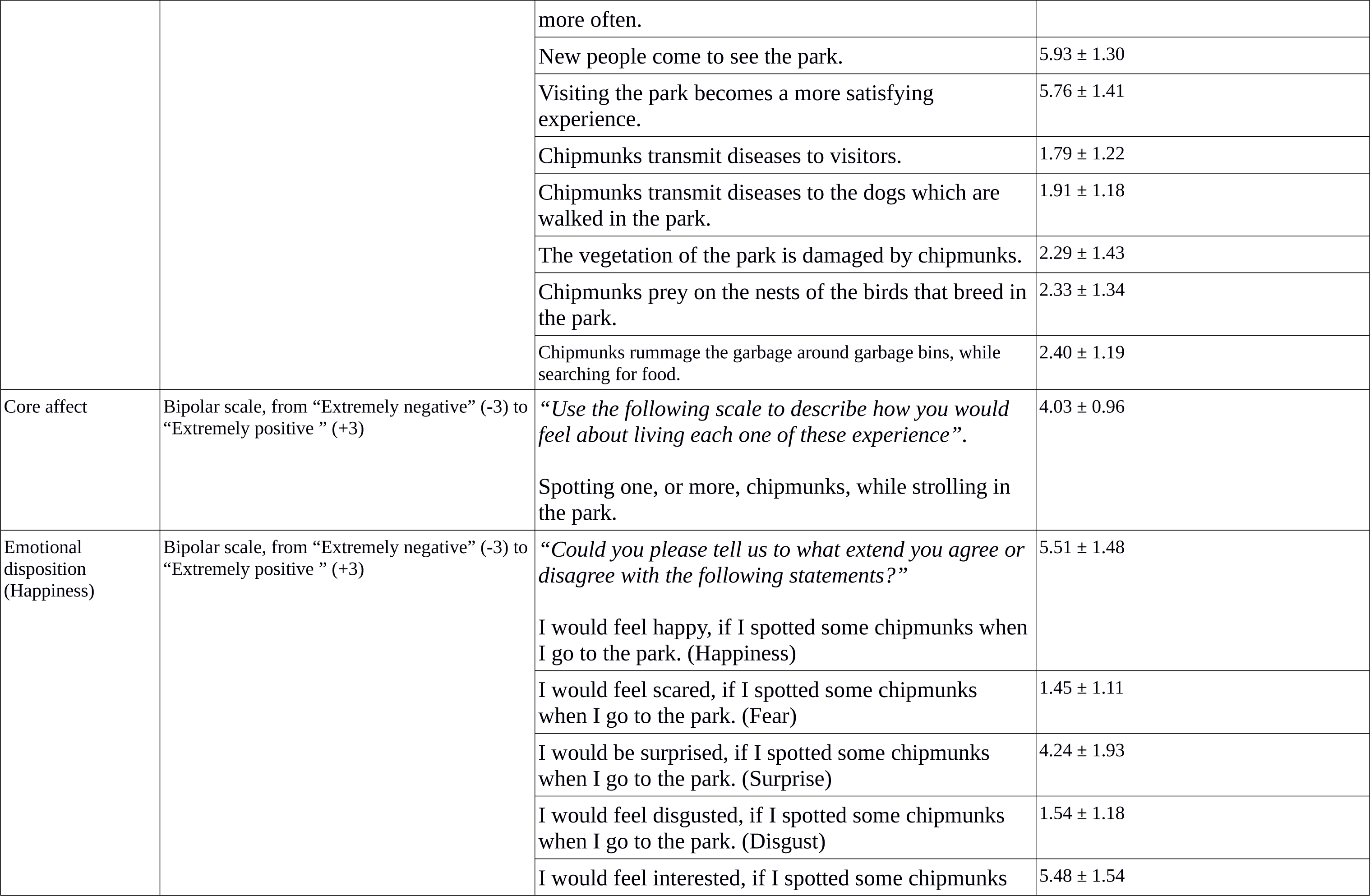
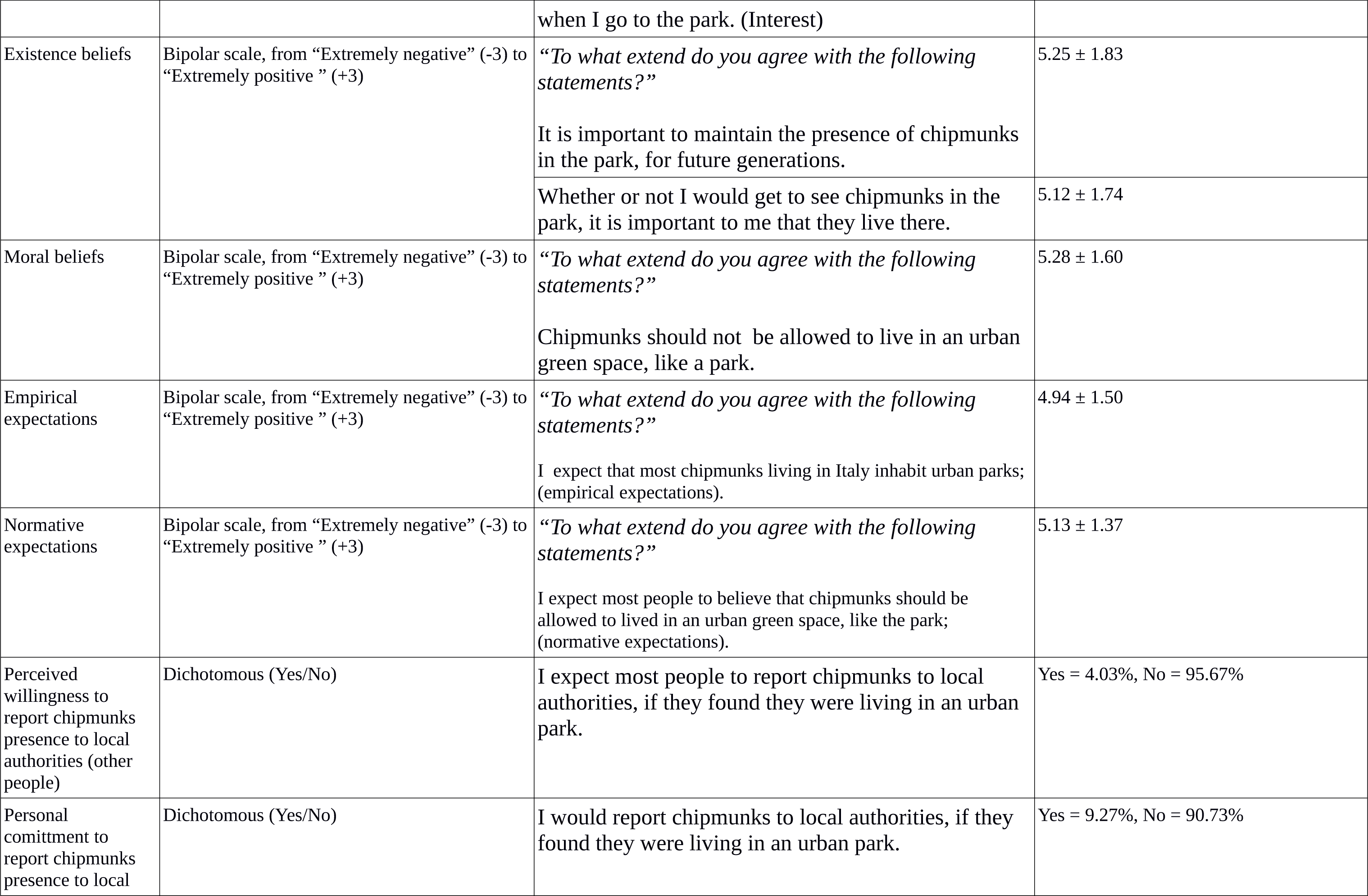
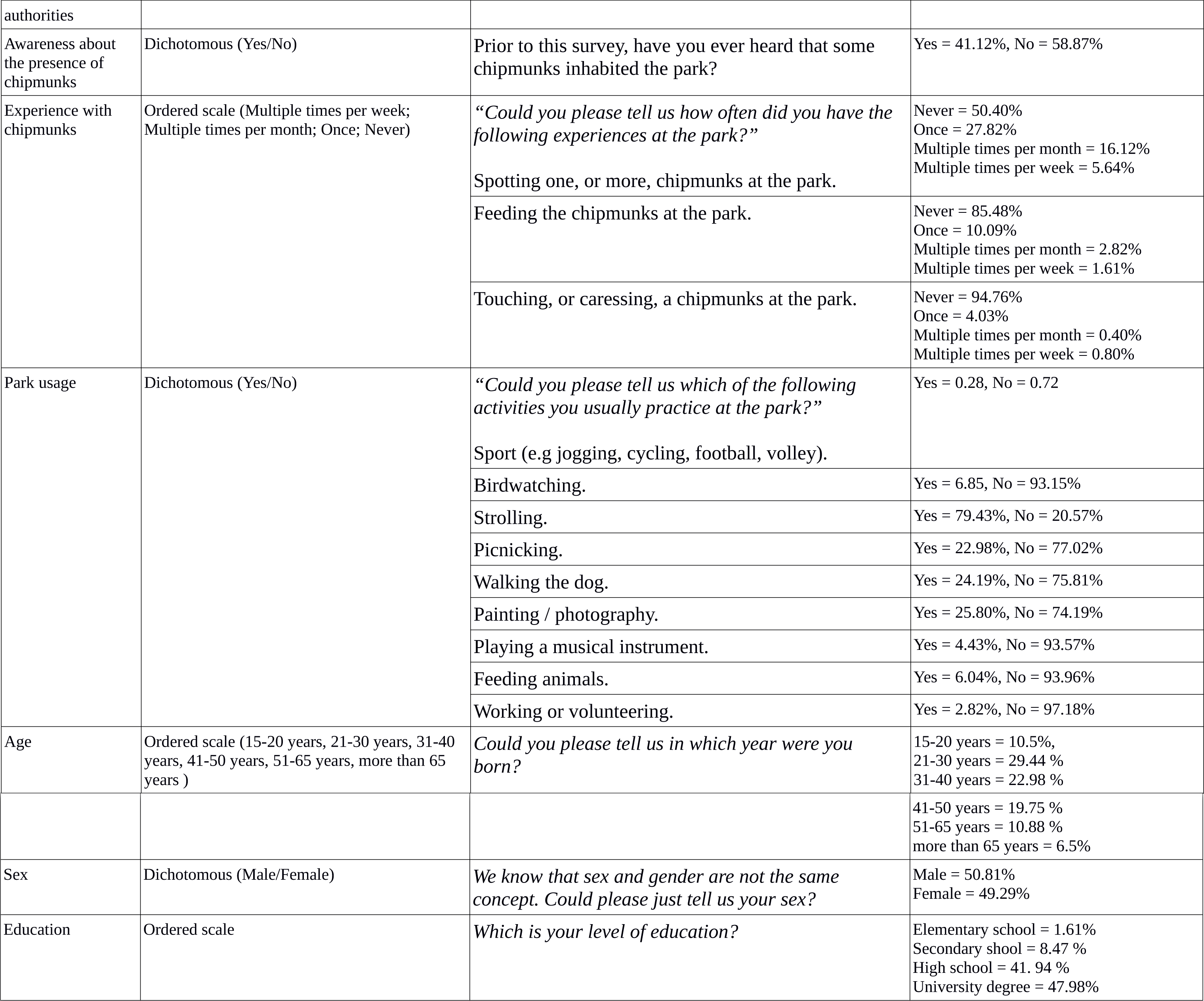
Questionnaire structure, item summary and completion rate.

We assessed the reliability of our attitudinal scale through McDonald’s Omega,^30^ and we tested for construct validity through Confirmatory Factor Analysis (CFA), with a Maximum Likelihood estimator with robust standard errors and a Satorra-Bentler scaled test statistic. Both indicator and latent variables were standardized and all the factor loadings were estimated. We adopted two correlated latent variables reflecting the strength of each beliefs ad the evaluation of its outcome and we also specified some residual correlations between each couple of items describing a specific impact. We selected the best subset of items and the best latent variable structure by comparing models through likelihood-ration testing and some fitness indexes. Attitudes were aggregated into a final score by summing the product of each couple of items.^23^

We segmented respondents on the basis of their attitudes, emotional dispositions, core affect and their moral, empirical and normative expectations about the presence of chipmunks in the park. Segmentation aimed to identify clusters of respondents who strongly supported chipmunk presence and could oppose their eradication. We tested for the presence of clusters in the data through the Hopkins index and we compared k-means, hierarchical and k-medoid cluster analysis^31^ to assess which one clustered observation the best.

We carried out Generalized Linear Modeling with a Gamma distribution of the error and an identity link to highlight differences in the two areas, in terms of attitudes scores, core affect, existence beliefs and moral, empirical and normative expectations about the presence of chipmunks in the park. In each model we included a dummy variable assessing whether respondents ecountered chipmunks at least once in their lifetime, and an interaction term, to account for the effect of past behavior and belief saliency.

Finally, we tested for chipmunk expansion outside of their introduction sites by means of geographic profiling (GP). GP is common in criminology, where the spatial locations of crimes are used to calculate the probability of occurrence of the offender’s residence for each point over a certain geographical area. GP outperforms classical measures of spatial tendency, and many ecologists found it good for tracing back the origin of individuals that could move across space.^32-35^ We adopted a Bayesian GP algorithm,^36^ requiring only the specification of a distribution parameter, indicating a plausible maximum extent to which individuals could move. Based on available evidence indicating that chipmunks usually disperse within a few hundred meters from their birthplace,^37-38^ we opted for a dispersal parameter of 1 km. We used available observations collected in Villa Ada, from 2011 to 2014 (n=26), and in Valeggio sul Mincio, from 1997 to 2018 (n=87), as the input for the GP algorithm. We did not use observations from Villa Doria-Pamphilii as the park is embedded in an urbanized matrix, which prevents chipmunks from dispersing around. It is important to note that we were not interested to identify where chipmunks were released, but to reconstruct a probabilistic profile for the origin of the observations: the inspection of its shape told us whether observed chipmunks came from disjoint hotspots, as expected for an expanding invasive alien species, or from a single one, like in the case of a species which is not expanding. Statistical analyses were carried out with the statistical software R (R Core Team 2018) and a detailed information about statistical analysis, altogether with a reproducible software code is available in the Supplementary Information (S2).

## 3 RESULTS AND DISCUSSION

Respondents had generally positive emotions towards chipmunks. Moreover, they generally agreed with the idea that the presence of chipmunks in the park was important for future generations and that it was important to have chipmunks living in the park even if one does not see them. Finally, most respondents deemed right and common for chipmunks to live in urban parks (Table 1). However, most respondents were not aware of the presence of chipmunks in the park where they were interviewed and about half of them had never observed these animals before. The proportion of respondents who had fed (14.6%) or touched (6.3%) chipmunks were even lower. Moreover, 14.11% of respondents reported to have observed chipmunks, despite they were not aware of their presence (Figure 1).

**Figure 1.**
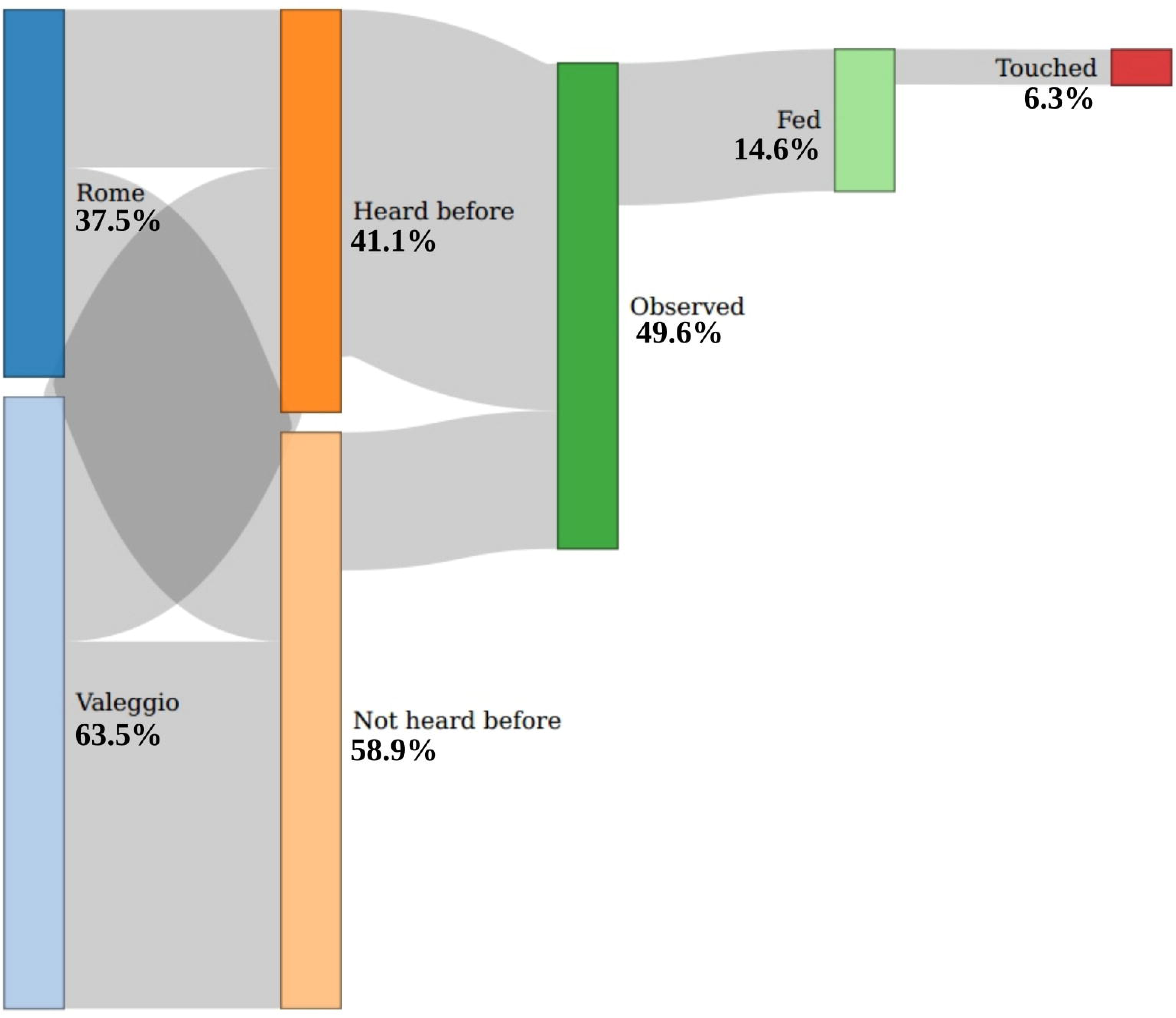
Sankey plot about visitors-chipmunks interactions.

CFA and McDonald’s Omega did not support an overall attitudinal construct, but they identified two separate groups of beliefs. The first one included items about the impact of chipmunks over the quality of recreation at the park: increasing the aesthetic appeal of the park, attracting new visitors and making visitors more prone to visit the park again. The second group included the potential impacts of chipmunks over human health: rummaging garbage from bins, transmitting disease to humans and to visitors’ dogs (Table 2; Table 3).

**Table 2.**
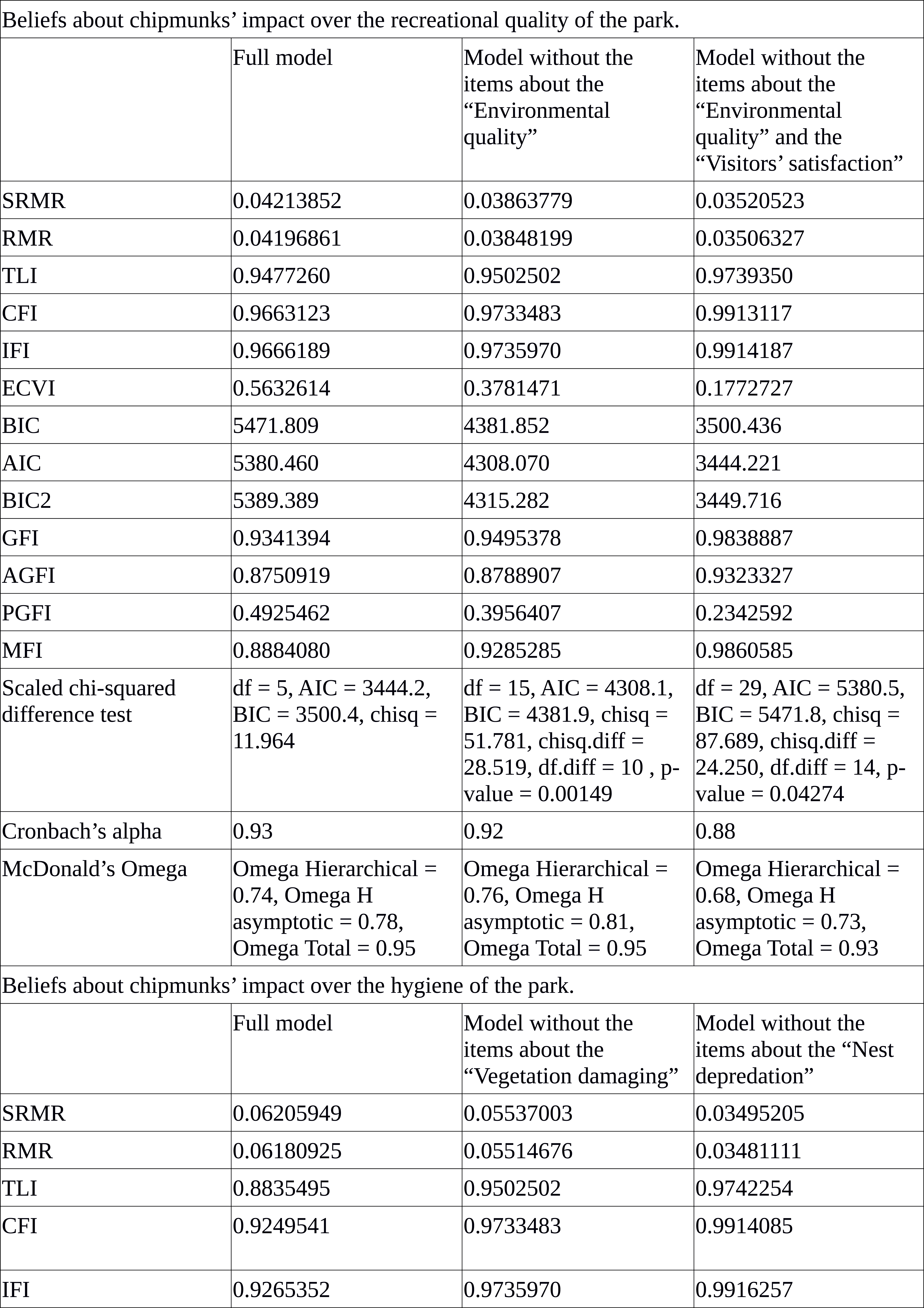
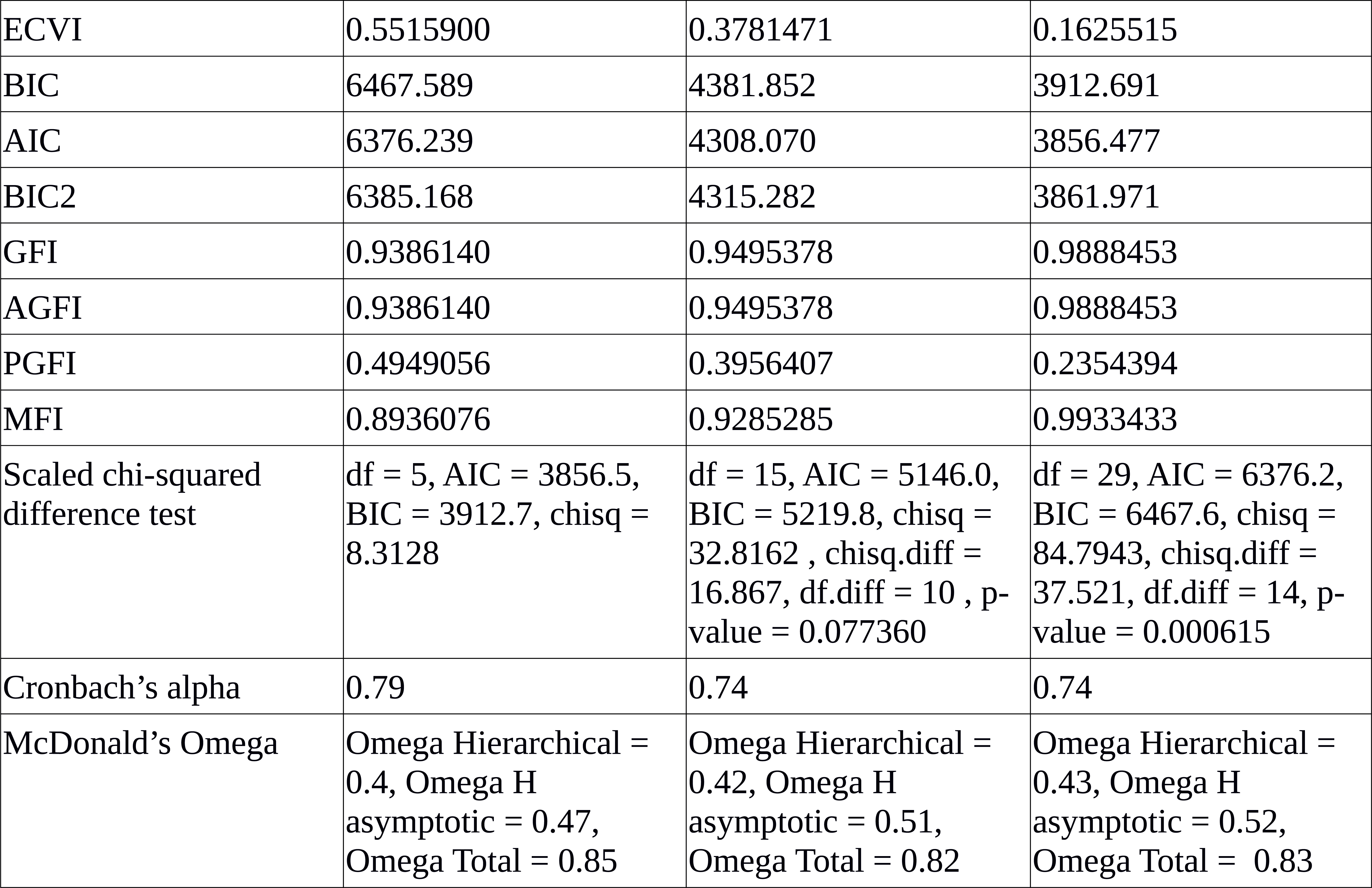
Confirmatory Factor Analysis, model comparison.

**Table 3.**
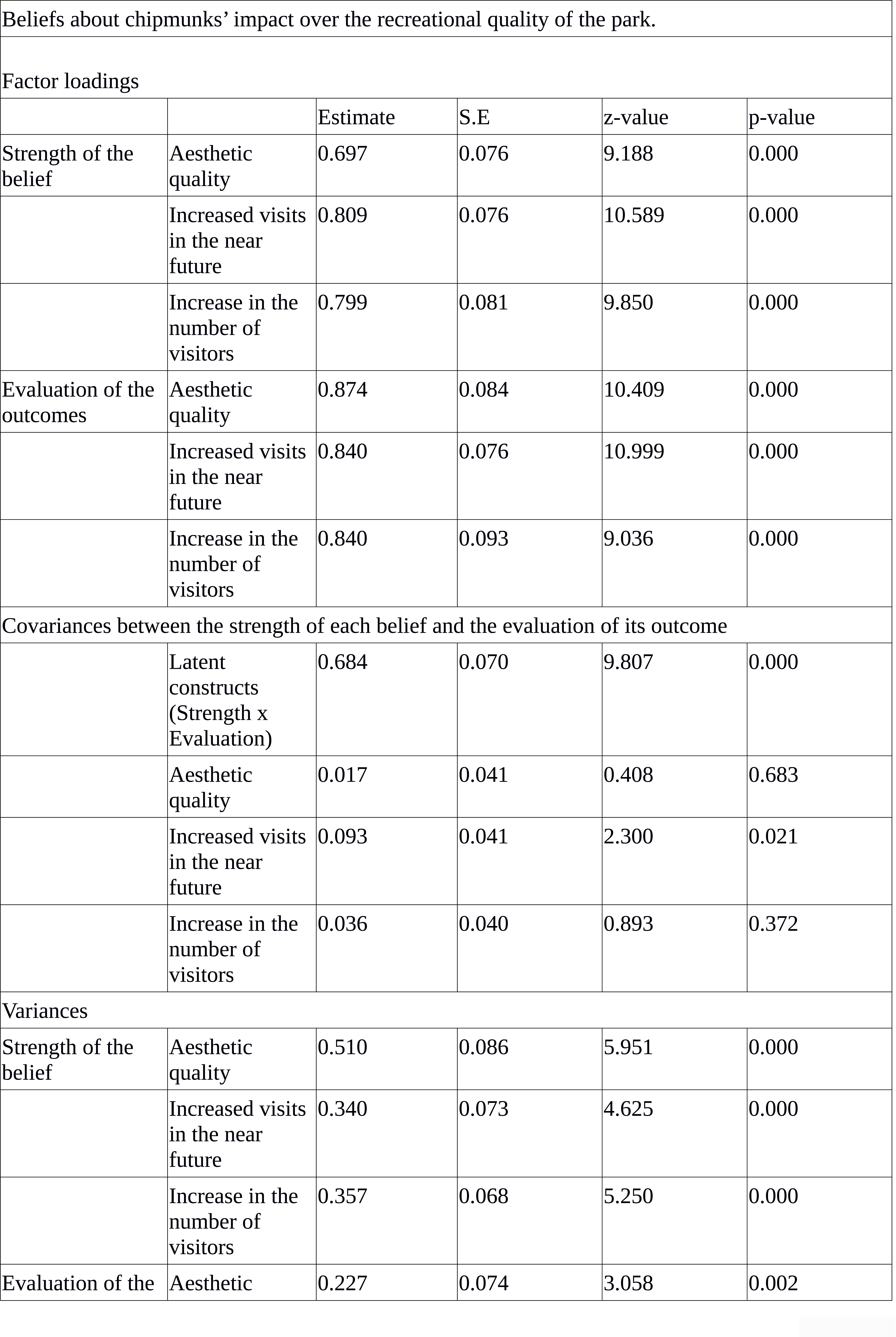
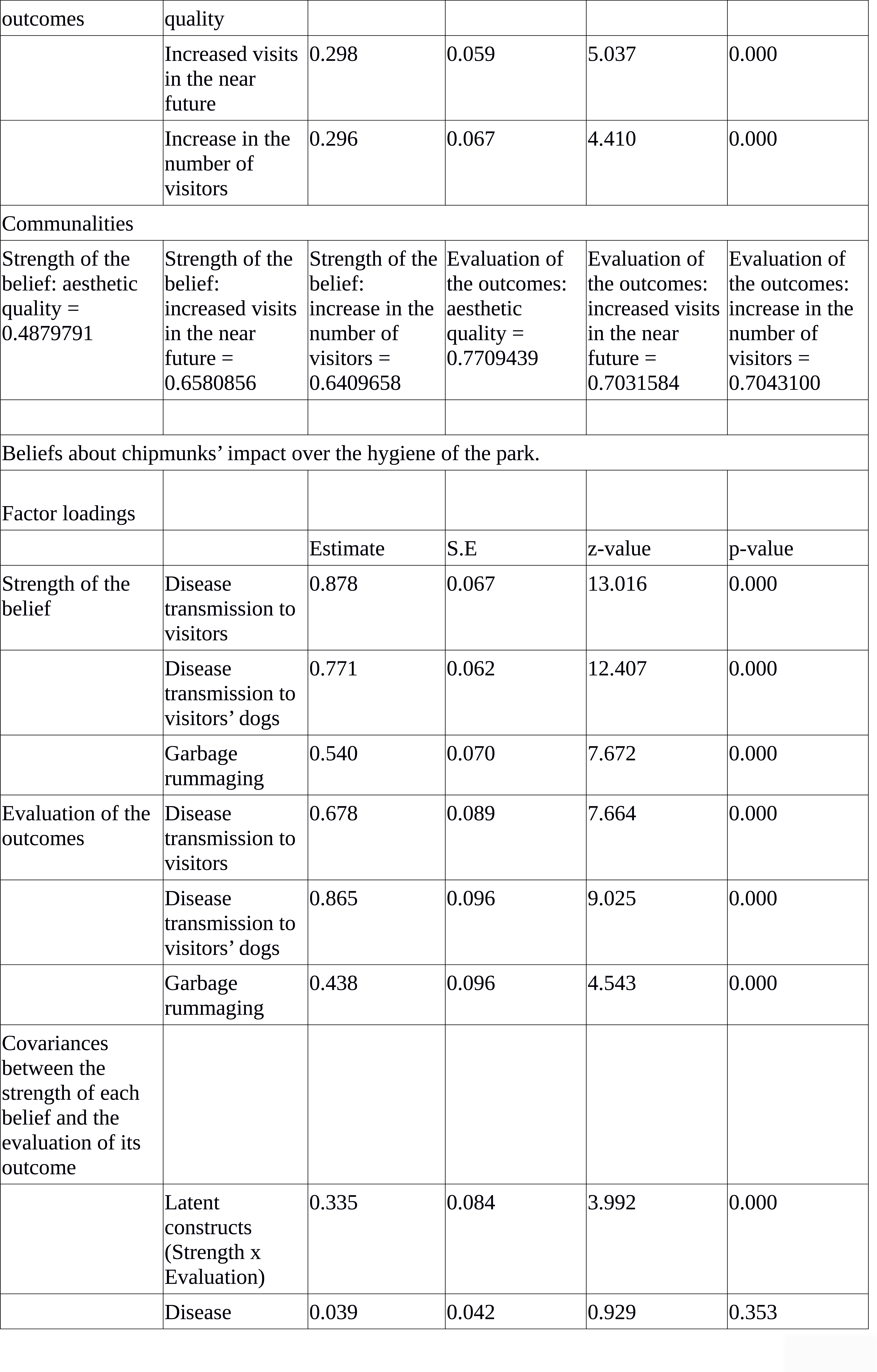
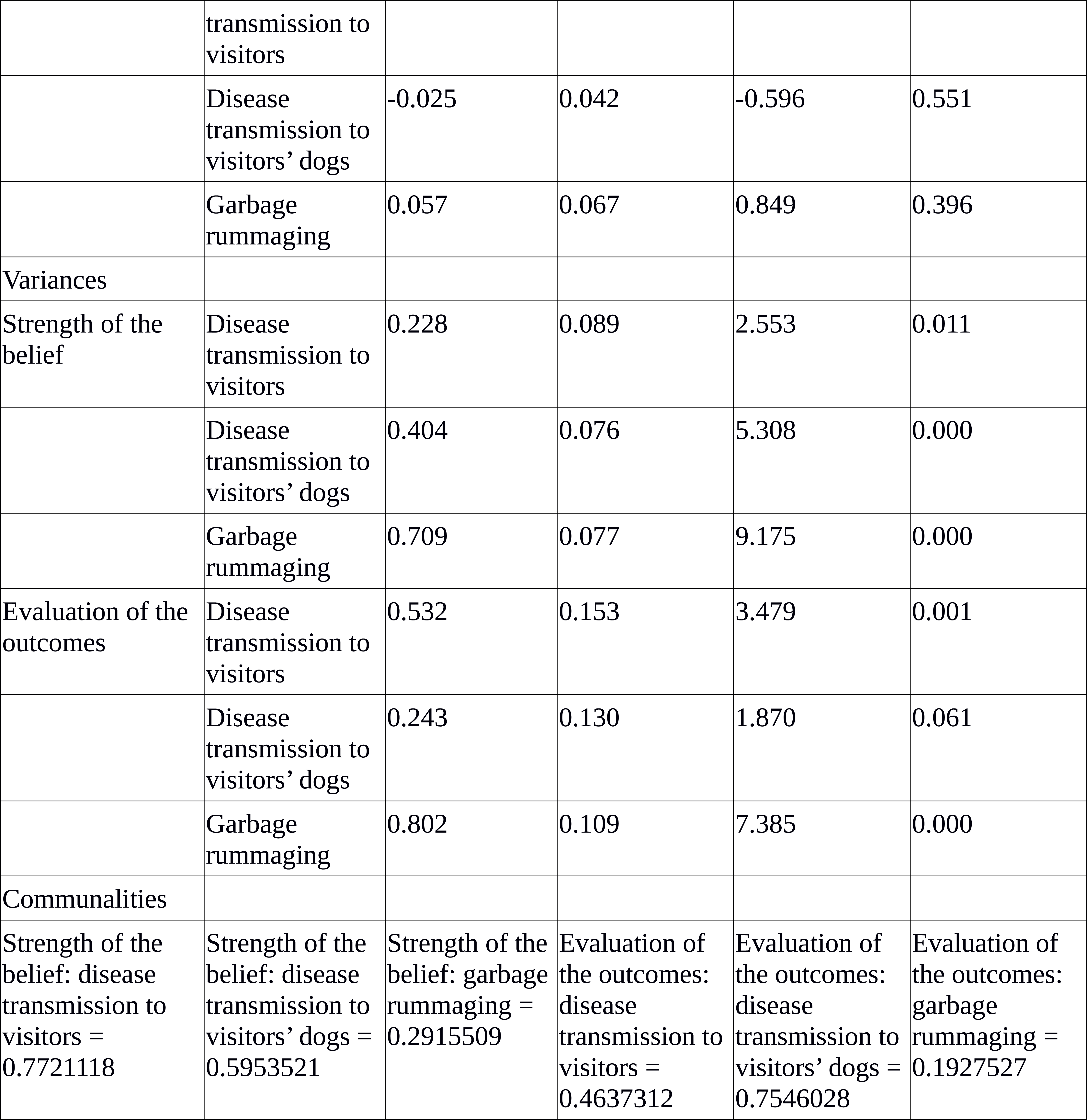
Confirmatory Factor Analysis: factor loadings, covariances, variances and communalities.

Hierarchical cluster analysis with Euclidean distance and a complete link indicated the presence of a small segment of respondents, characterized by negative attitudes about chipmunk impacts over human health, fear and disgust towards chipmunks (Figure 2). These could be people who are scared or disgusted by rodents and concerned about their impact over hygiene, two aspects that are often related.^39-41^

**Figure 2.**
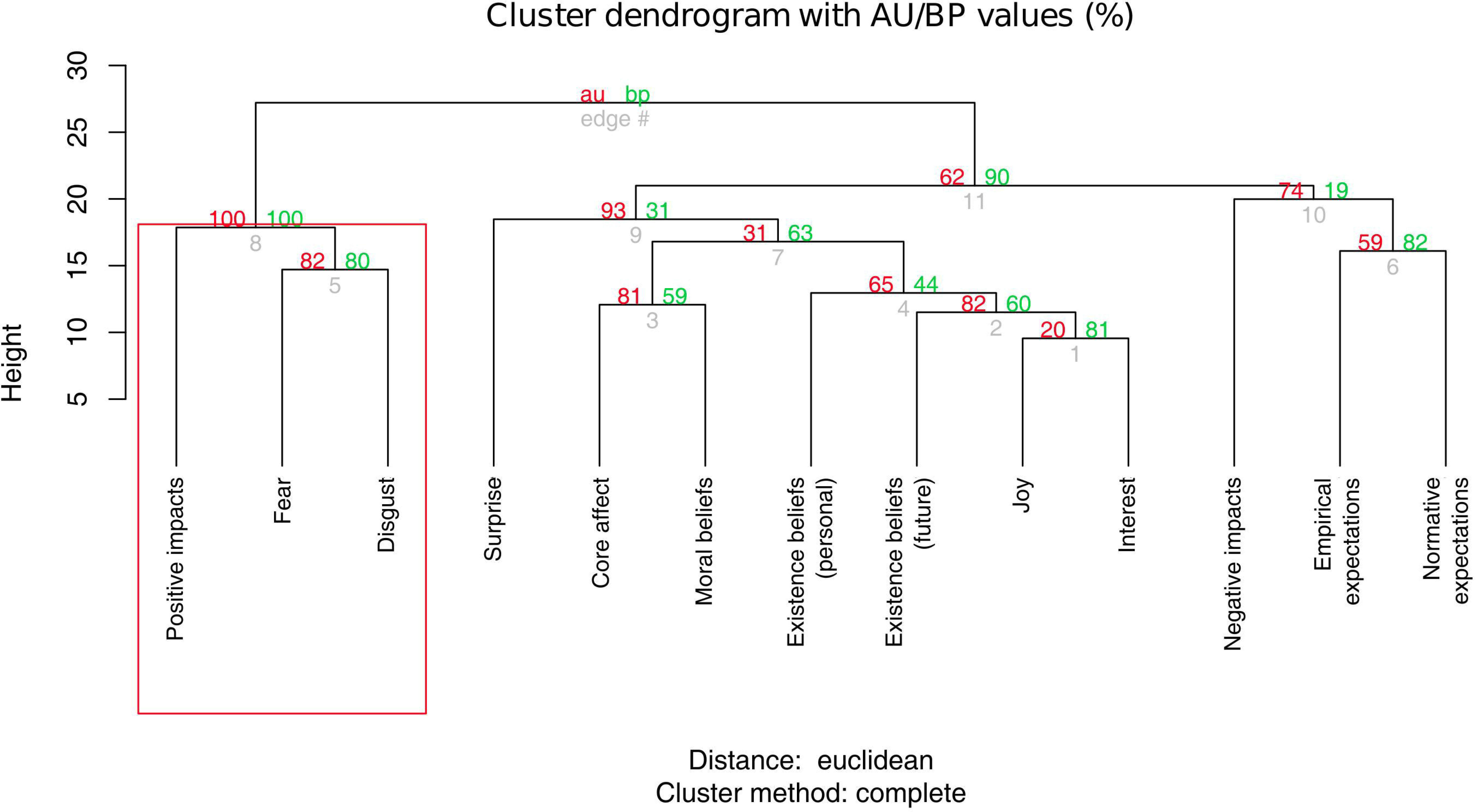
Dendrogram of the hierarchical cluster analysis.

The two sites differed only in respondents’ score about the positive impact of chipmunks over the recreational experience, with Rome having slightly higher scores. Visitors in Rome also agreed slightly more with the idea that most people deemed appropriate for chipmunks to live in urban parks in Italy (Figure 3; Table 4).

**Table 4.**
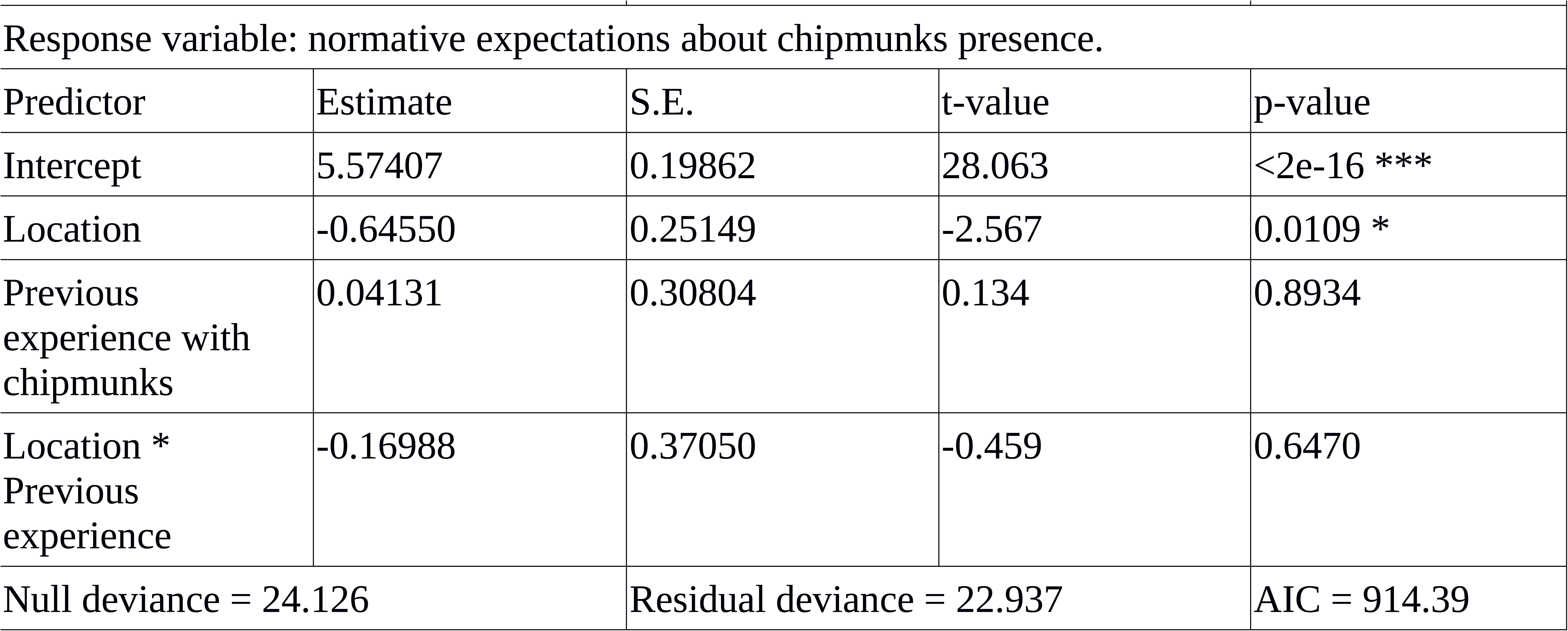
Ouput of the Generalized Linear Model. Normative expectations were measured on a rescaled biploar Likert scales, ranging from 1 to 7.

**Figure 3.**
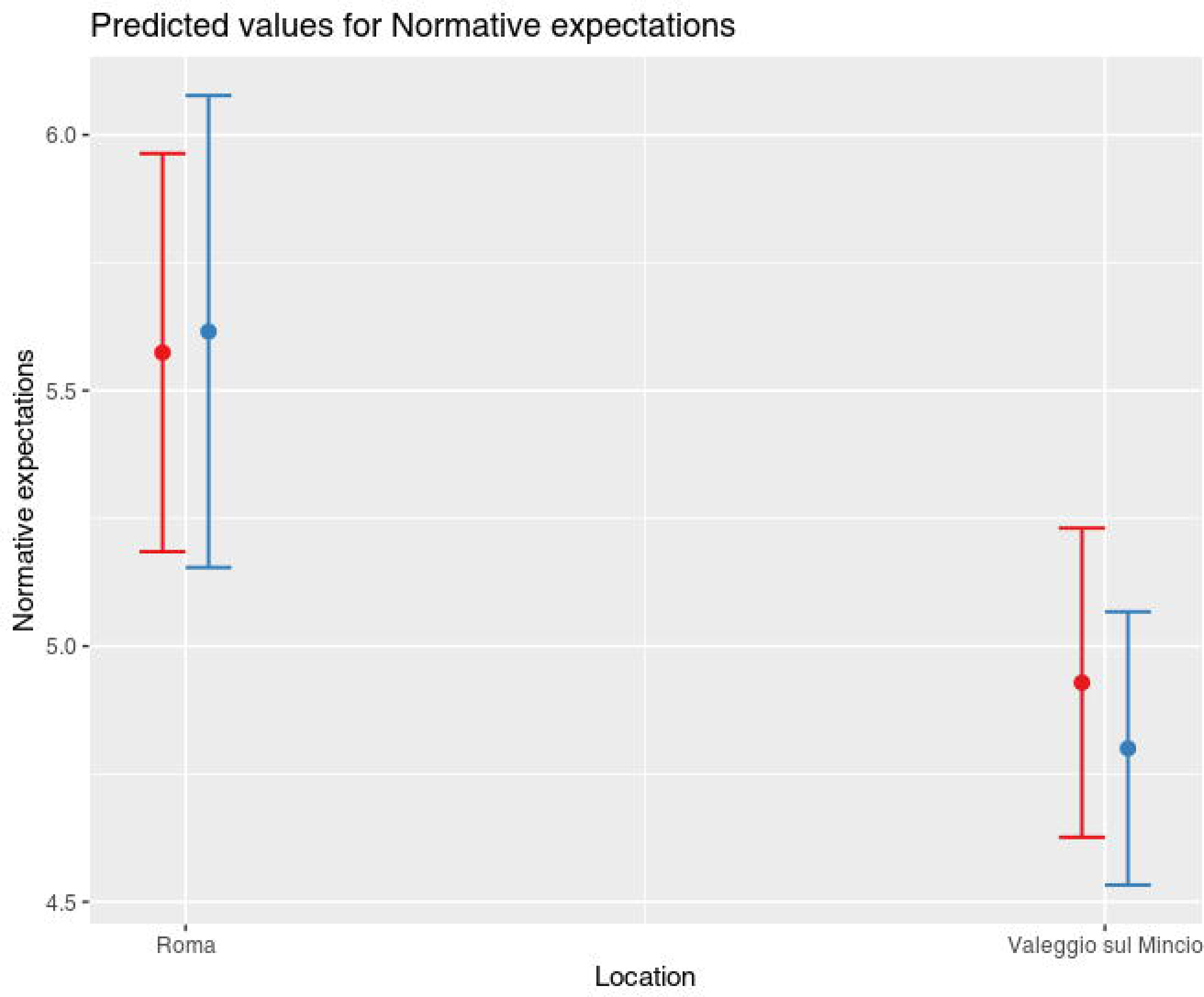
Marginal effects of the GLM, based on a Gamma distribution of the error and an identity link: average scores of normative expectations in the two areas, both for respondents who had observed chipmunks (blue) and for respondents who didn’t (red).

Overall, these findings indicate that visitors regard chipmunks positively. Apart from a small cluster of respondents, most of them have positive emotions towards chipmunks, deem appropriate the fact that they live in an urban park and value their presence as something having an intrinsic value. However, this positive perception probably stems from a more general, favorable, disposition towards the presence of wildlife at urban areas. Respondents do not have a coherent system of attitudes about the presence of chipmunks, probably because their real interactions were limited: attitudes are shaped and reinforced by our everyday experience with a certain issue, that make it salient for ourselves.^23,42^ On the other hand, visitors had stable beliefs about those impacts of chipmunks that could affect their recreational experience at the park, as well as fears about those impacts that could undermine hygiene. These two sub-dimensions probably indicate that visitors’ beliefs are embedded in broader belief networks encompassing different, and more salient, topics.^43^ For example, our respondents could have stable belief networks diseases, and they could have tied to them some of their beliefs about chipmunks. Framing experiments, where participants are primed to think about some precise topics and where the effect of this priming over beliefs is measured^44^ might be a valuable tool to better investigate how human-wildlife interactions are embedded into broader nomological networks, and influenced by beliefs about relevant social issues. Framing experiments could also be used to test for attitude certainty and strength.^45,46^

The idea that respondents’ attitudes were not grounded into experience is reinforced by the limited interactions between visitors and chipmunks: approximately, only half visitors observed chipmunks in the park, 15% of them fed chipmunks and only about 6% of them reported to have touched a chipmunk. Moreover, some visitors who observed chipmunks were not aware of their stable presence in the park: our questionnaire was arguably the first time they were introduced to this aspect. These superficial interactions are also reflected by the low differences between the two areas. Respondents in Rome and in Valeggio sul Mincio had similar scores for almost all the psychological antecedents of their interactions with chipmunks. They showed minor differences only in their beliefs about the impact of chipmunks over the recreational experience, and in their normative beliefs about the presence of chipmunks in an urban green area. Moreover, hierarchical cluster analysis did not divide respondents into meaningful segments, on the basis of their attitudes, emotions, existence values and social norms towards chipmunks and it did not identify any group of strong supporters of chipmunks.

Taken together, these three points are important for the future management of chipmunks in the study area. Attitudes are an antecedent of human behavior and often a good barometer to forecast an eventual opposition to the management of invasive native^23^ and introduced wildlife.^47^ As visitors do not have stable attitudes and no segments of highly motivated visitors exist, it is reasonable to say that an eradication campaign would not face any strong opposition from local visitors. Chipmunks at the two sites do not seem to be an iconic species yet like the gray squirrel in many urban parks if the UK.^48^ Their interactions with visitors, especially those creating emotional bindings, like feeding, are still limited. However, considered that respondents regard chipmunks favorably and that they value their presence as a legacy for the younger generations, we believe that eradication initiatives should be coupled with an adequate communication strategy, to avoid polarization and the ‘backfire effect’. Considered that respondents from the two sites did not show any particular difference, communication actions might be similar for Rome and Valeggio sul Mincio.

Geographic profiling confirmed that invasive chipmunks disperse less than 500 m from the place where they are born. However, while chipmunk observations in Rome come from a single source, observations in Valeggio sul Mincio are likely to have involved individuals coming from two distinct spatial cores (Figure 4). One of these two cores was found to be outside of the boundaries of the urban park where the species was introduced: although slowly, chipmunks are expanding outside Sigurtà Garden Park, their introduction site in Valeggio sul Mincio. As chipmunks increase their density we expect them to continue their expansion outside of the park. This perspective is not encouraging, because Valeggio sul Mincio is surrounded by a countryside environment and cultivations, that might promote chipmunks dispersal at the landscape scale^20^ and maybe even their role as a pest species damaging crops, in the near future. Considered that chipmunks are still distributed over a relatively small area, but there are large areas in Italy suitable for the species ^49,50^, we recommend their quick removal, as it will be easy and cost-effective.

**Figure 4.**
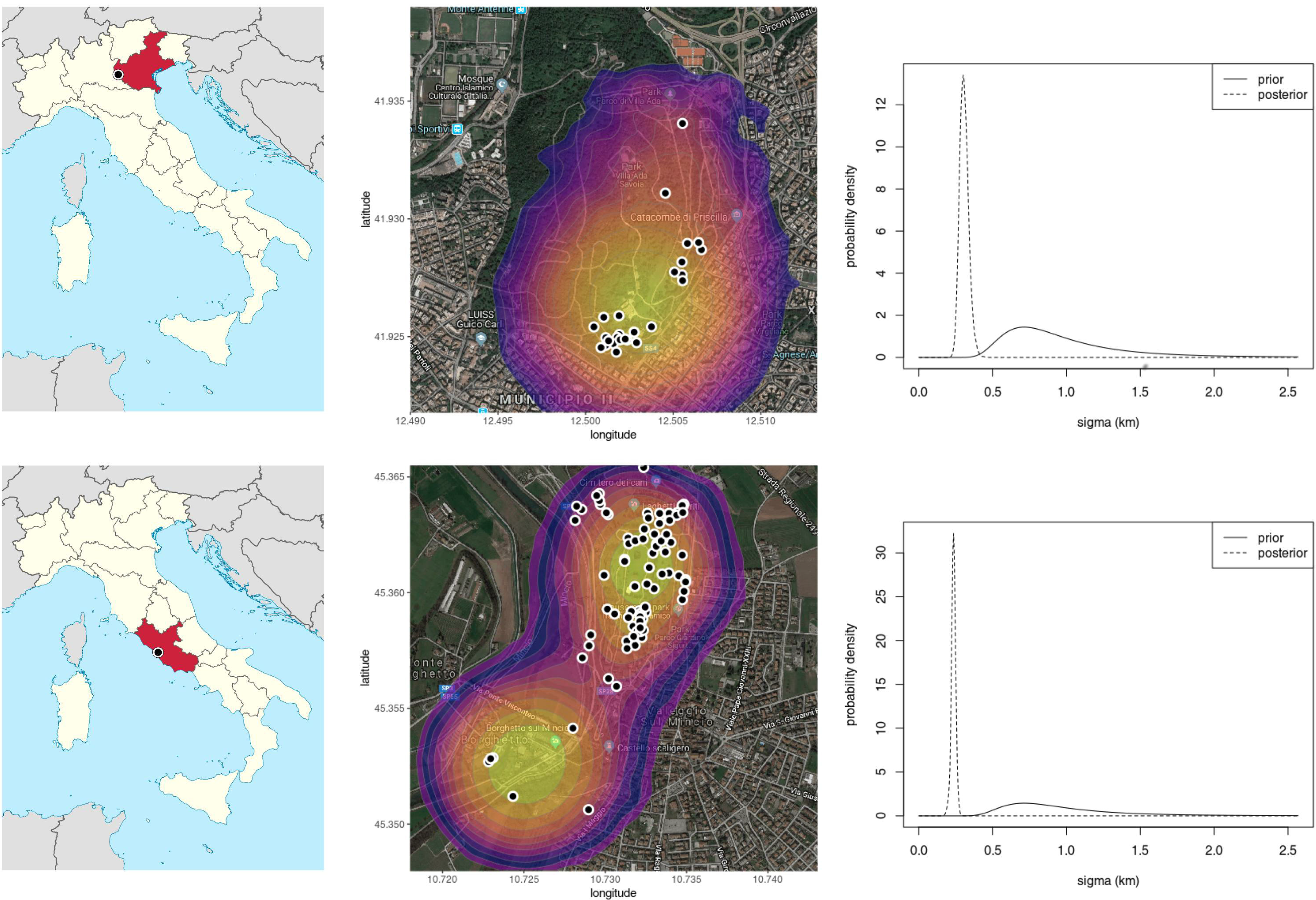
Location of the three study sites (left), heatmap with the posterior probabilities of the origins of observed tamias (center) and posterior probability of the dispersal parameter (right).

## 4 CONCLUSION

In this research, we showed how spatially-explicit data about a biological invasion and survey data about its social perception can inform decision makers about the feasibility, and the urgency, of management actions. Geographic profiling can be used not only to identify introduction, or to locate dens of invasive pests but also to signal the emergence of source-sink systems. These systems indicate the end of an early invasion stage and the spatial expansion of the invasive alien species, often due to their numerical increase, which can make eradication or control harder and expensive. Moreover, structured surveys could inform conservationists about the interactions between stakeholders and biological invaders, altogether with their social perception.

Our findings indicate that visitors still have limited interactions with invasive chipmunks, at the urban parks in Italy where they have been introduced. They do not have stable attitudes, and there seems not to be any group of visitors who regard chipmunks as an iconic species. Perspectives about chipmunks are positive, but probably weak. At the same time, chipmunks are expanding outside of their introduction site in Northern Italy. We deem that initiatives aimed at removing chipmunks are still feasible, if properly planned, and urgent. Postponing any management intervention could complicate eradications, both for the spatial spread of the species in one of the two areas and for the risk of change in visitors’ attitude due to a greater confidence with chipmunks that have become more abundant.

## Supporting information

Dataset from attitude scaling pilot

Reproducible R script

Dataset used in the analyses

## ACKNOWLEDGMENTS

The “U.O. Manutenzione e Valorizzazione del Verde Urbano” office of the Municipality of Roma provided us with permits to set the survey within urban parks in Rome. Authors would like to thank Davide Sogliani, who kindly helped in the fieldwork, and Giulia Benassi, who provided them with data from previous surveys about chipmunks presence in Rome.

