## Supplementary material for "Invasive Siberian chipmunks *Eutamias sibiricus* in Italy: a socio-ecological analysis indicates that they could, and should, be removed": Reproducible R script: S2.R

```
#Loading packages
{
require(tidyverse)
require(plyr)
require(lubridate)
require (devtools)
require(tidyselect)
require(semTools)
require(psy)
require(psych)
require(Amelia)
#require(mice)
require(semPlot)
require(ggeffects)
#require(gridExtra)
#require(data.table)
#require(sjstats)
#require(sjlabelled)
#require(sjmisc)
require(sjPlot)
require(cluster)
require(factoextra)
require(fpc)
#require(NbClust)
#require(clValid)
#require(clustertend)
#require(pvclust)
require(RgeoProfile)
require(devtools)
require(ggmap)
register_google(key ="AIzaSyDV9QP7CdmQ3YApschivng59DoRY3rs8Rg")
require(maps)
require(maptools)
require(sp)
require(rgdal)
#require(xlsxjars)
#require(xlsx)
require(raster)
#require(stringr)
require(foreign)
#require(networkD3)
require(clustertend)
require(clValid)
require(pvclust)
}

################################################################
#Dataset
################################################################
#Loading the dataset
{
df <- read.csv(file="~/Documents/Jacopo/Research_Projects/Ongoing/Tamias/Data/Dataset_tamias.csv")
head(df)
nrow(df)
df <- df[-249,]
names(df)
table(df$Location)
}

#Renaming factors and levels
{
df$ID <- rownames(df)
df$Location <- as.character(factor(df$Location))

df$Presence_before <- factor(df$Presence_before)

df$Education <- plyr::revalue(df$Education, c("Il diploma di scuola superiore"="High_school", 
  "Il diploma di scuola superiore, La laurea"="Degree", "La laurea"="Degree",
  "La licenza elementare"="Elementary", "La licenza elementare, La licenza media"="Secondary_school",
  "La licenza elementare, La licenza media, Il diploma di scuola superiore"="High_school",
  "La licenza elementare, La licenza media, Il diploma di scuola superiore, La laurea"="Degree",
  "La licenza media"="Secondary_school", "La licenza media, Il diploma di scuola superiore"="High_school"))
df$Education <- factor(df$Education, levels=c("Elementary", "Secondary_school", "High_school", "Degree"))

df[which(df$Frequency_visit==""), "Frequency_visit"] <- NA 
df$Frequency_visit <- factor(df$Frequency_visit)
df$Frequency_visit <- factor(df$Frequency_visit, levels=c("Mai, oggi è la prima volta.",
                            "Ci sono venuto solo un' altra volta.", "Più volte in un anno.",
                            "Più volte al mese.", "Più volte alla settimana"))
levels(df$Frequency_visit)
df$Frequency_visit <- plyr::revalue(df$Frequency_visit, c("Mai, oggi è la prima volta."="First time",
                      "Ci sono venuto solo un' altra volta."="Second time", 
                      "Più volte in un anno."="Multiple times per year", 
                      "Più volte al mese."="Multiple times per month",
                      "Più volte alla settimana"="Multiple times per week"))
levels(df$Frequency_visit)

levels(df$Squirrel_watching)
df[which(df$Squirrel_watching==""), "Squirrel_watching"] <- NA 
df$Squirrel_watching <- factor(df$Squirrel_watching)
df$Squirrel_watching <- plyr::revalue(df$Squirrel_watching, c("Mai"="Never", 
                        "E' successo una volta sola"="Once", 
                        "Più volte al mese"="Multiple times per month",
                        "Più volte a settimana"="Multiple times per week"))

levels(df$Squirrel_feeding)
df[which(df$Squirrel_feeding==""), "Squirrel_feeding"] <- NA 
df$Squirrel_feeding <- factor(df$Squirrel_feeding)
df$Squirrel_feeding <- plyr::revalue(df$Squirrel_feeding, c("Mai"="Never", 
                        "E' successo una volta sola"="Once", 
                        "Più volte al mese"="Multiple times per month",
                        "Più volte a settimana"="Multiple times per week"))

levels(df$Squirrel_touching)
df[which(df$Squirrel_touching==""), "Squirrel_touching"] <- NA 
df$Squirrel_touching <- factor(df$Squirrel_touching)
df$Squirrel_touching <- plyr::revalue(df$Squirrel_touching, c("Mai"="Never", 
                       "E' successo una volta sola"="Once", 
                       "Più volte al mese"="Multiple times per month",
                       "Più volte a settimana"="Multiple times per week"))

df[which(df$Joy==""), "Joy"] <- NA 
df$Joy <- factor(df$Joy)
table(df$Joy)
df$Joy <- plyr::revalue(df$Joy, c("Molto in disaccordo"="Strongly disagree",
                                  "In disaccordo"="Disagree",
                                  "Leggermente in disaccordo"="Mildly disagree",
                                  "Non saprei"="I don't know",
                                  "Leggermente d'accordo"="Mildly agree",
                                  "D'accordo"="Agree",
                                  "Molto d'accordo"="Strongly agree"))
df$Joy <- factor(df$Joy, levels=c("Strongly disagree", "Disagree", "Mildly disagree",
                                  "I don't know", "Mildly agree", "Agree", "Strongly agree"))

df[which(df$Fear==""), "Fear"] <- NA 
df$Fear <- factor(df$Fear)
df$Fear <- plyr::revalue(df$Fear, c("Molto in disaccordo"="Strongly disagree",
                                  "In disaccordo"="Disagree",
                                  "Leggermente in disaccordo"="Mildly disagree",
                                  "Non saprei"="I don't know",
                                  "Leggermente d'accordo"="Mildly agree",
                                  "D'accordo"="Agree",
                                  "Molto d'accordo"="Strongly agree"))
df$Fear <- factor(df$Fear, levels=c("Strongly disagree", "Disagree", "Mildly disagree",
                                  "I don't know", "Mildly agree", "Agree", "Strongly agree"))

df[which(df$Surprise==""), "Surprise"] <- NA 
df$Surprise <- factor(df$Surprise)
df$Surprise <- plyr::revalue(df$Surprise, c("Molto in disaccordo"="Strongly disagree",
                     "In disaccordo"="Disagree",
                     "Leggermente in disaccordo"="Mildly disagree",
                     "Non saprei"="I don't know",
                     "Leggermente d'accordo"="Mildly agree",
                     "D'accordo"="Agree",
                     "Molto d'accordo"="Strongly agree"))
df$Surprise <- factor(df$Surprise, levels=c("Strongly disagree", "Disagree", "Mildly disagree",
                                    "I don't know", "Mildly agree", "Agree", "Strongly agree"))

df[which(df$Disgust==""), "Disgust"] <- NA 
df$Disgust <- factor(df$Disgust)
df$Disgust <- plyr::revalue(df$Disgust, c("Molto in disaccordo"="Strongly disagree",
                    "In disaccordo"="Disagree",
                    "Leggermente in disaccordo"="Mildly disagree",
                    "Non saprei"="I don't know",
                    "Leggermente d'accordo"="Mildly agree",
                    "D'accordo"="Agree",
                                            "Molto d'accordo"="Strongly agree"))
df$Disgust <- factor(df$Disgust, levels=c("Strongly disagree", "Disagree", "Mildly disagree",
                                            "I don't know", "Mildly agree", "Agree", "Strongly agree"))

df[which(df$Interest==""), "Interest"] <- NA 
df$Interest <- factor(df$Interest)
df$Interest <- plyr::revalue(df$Interest, c("Molto in disaccordo"="Strongly disagree",
                     "In disaccordo"="Disagree",
                     "Leggermente in disaccordo"="Mildly disagree",
                     "Non saprei"="I don't know",
                     "Leggermente d'accordo"="Mildly agree",
                     "D'accordo"="Agree",
                     "Molto d'accordo"="Strongly agree"))
df$Interest <- factor(df$Interest, levels=c("Strongly disagree", "Disagree", "Mildly disagree",
                                          "I don't know", "Mildly agree", "Agree", "Strongly agree"))
df$Squirrel_exp <- NA
df[which(df$Squirrel_feeding=="Never"&df$Squirrel_touching=="Never"&df$Squirrel_watching=="Never"),
   "Squirrel_exp"] <- 0
df[which(is.na(df$Squirrel_exp)),"Squirrel_exp"]<-1
prop.table(table(df$Squirrel_exp))

df[which(df$MoralBel==""), "MoralBel"] <- NA 
df$MoralBel <- factor(df$MoralBel)
levels(df$MoralBel)
df$MoralBel <- plyr::revalue(df$MoralBel, c("Profondamente sbagliato"="Absolutely inappropriate",
               "Sbagliato"="Inappropriate", "Leggermente sbagliato"="Slightly inappropriate",
               "Né giusto, né sbagliato. Non saprei." = "Neither wrong, nor right. I don't know",
               "Leggermente giusto"="Basically right", "Giusto"="Right", 
               "Profondamente giusto"="Absolutely right"))
df$MoralBel <- factor(df$MoralBel, levels=c("Absolutely inappropriate", "Inappropriate",
               "Slightly inappropriate", "Neither wrong, nor right. I don't know",
               "Basically right", "Right", "Absolutely right"))

table(df$EmpExp_Presence)
df[which(df$EmpExp_Presence==""), "EmpExp_Presence"] <- NA 
df$EmpExp_Presence <- factor(df$EmpExp_Presence)
levels(df$EmpExp_Presence)
df$EmpExp_Presence <- plyr::revalue(df$EmpExp_Presence, c("Molto in disaccordo"="Strongly disagree",
                    "In disaccordo"="Disagree",
                    "Leggermente in disaccordo"="Mildly disagree",
                    "Non saprei"="I don't know",
                    "Leggermente d'accordo"="Mildly agree",
                    "D'accordo"="Agree","Molto d'accordo"="Strongly agree"))
df$EmpExp_Presence <- factor(df$EmpExp_Presence, levels=c("Strongly disagree", "Disagree", "Mildly disagree",
                      "I don't know", "Mildly agree", "Agree", "Strongly agree"))

df$NormExp_Acceptable
table(df$NormExp_Acceptable)
df[which(df$NormExp_Acceptable==""), "NormExp_Acceptable"] <- NA 
df$NormExp_Acceptable <- factor(df$NormExp_Acceptable)
levels(df$NormExp_Acceptable)
df$NormExp_Acceptable <- plyr::revalue(df$NormExp_Acceptable, c("Molto in disaccordo"="Strongly disagree",
                          "In disaccordo"="Disagree",
                          "Leggermente in disaccordo"="Mildly disagree",
                          "Non saprei"="I don't know",
                          "Leggermente d'accordo"="Mildly agree",
                          "D'accordo"="Agree","Molto d'accordo"="Strongly agree"))
df$NormExp_Acceptable <- factor(df$NormExp_Acceptable, levels=c("Strongly disagree", "Disagree", "Mildly disagree",
                                                          "I don't know", "Mildly agree", "Agree", "Strongly agree"))

df[which(df$EmpExp_Denounce==""), "EmpExp_Denounce"] <- NA 
df[which(df$PersonalCommitment_Denounce==""), "PersonalCommitment_Denounce"] <- NA 

levels(df$ExBel_future)
df[which(df$ExBel_future==""), "ExBel_future"] <- NA 
df$ExBel_future <- factor(df$ExBel_future)
levels(df$ExBel_future)
df$ExBel_future <- plyr::revalue(df$ExBel_future, c("Molto in disaccordo"="Strongly disagree",
                         "In disaccordo"="Disagree",
                         "Leggermente in disaccordo"="Mildly disagree",
                         "Non saprei"="I don't know",
                         "Leggermente d'accordo"="Mildly agree",
                         "D'accordo"="Agree","Molto d'accordo"="Strongly agree"))
df$ExBel_future <- factor(df$ExBel_future, levels=c("Strongly disagree", "Disagree", 
                          "Mildly disagree","I don't know", "Mildly agree", "Agree",
                          "Strongly agree"))
table(df$ExBel_future)

levels(df$ExBel_personal)
df[which(df$ExBel_personal==""), "ExBel_personal"] <- NA 
df$ExBel_personal <- factor(df$ExBel_personal)
levels(df$ExBel_personal)
df$ExBel_personal <- plyr::revalue(df$ExBel_personal, c("Molto in disaccordo"="Strongly disagree",
                    "In disaccordo"="Disagree",
                    "Leggermente in disaccordo"="Mildly disagree",
                    "Non saprei"="I don't know",
                    "Leggermente d'accordo"="Mildly agree",
                    "D'accordo"="Agree","Molto d'accordo"="Strongly agree"))
df$ExBel_personal <- factor(df$ExBel_personal, levels=c("Strongly disagree", "Disagree", 
                    "Mildly disagree","I don't know", "Mildly agree", "Agree", "Strongly agree"))
table(df$ExBel_personal)

levels(df$Age)
df[which(df$Age==""), "Age"] <- NA 
df$Age <- factor(df$Age, labels=c("15-20", "21-30", "31-40", "41-50", "51-65", "Over 65"))

df$Squirrel_watching <- factor(df$Squirrel_watching, levels=c("Never", "Once",
                              "Multiple times per month", "Multiple times per week"))
df$Squirrel_feeding <- factor(df$Squirrel_feeding, levels=c("Never", "Once",
                              "Multiple times per month", "Multiple times per week"))
df$Squirrel_touching <- factor(df$Squirrel_touching, levels=c("Never", "Once",
                              "Multiple times per month", "Multiple times per week"))

names(df)
df[,c(1:10)==""]
df[,1:10][df[,1:10] == "" ] <- NA
df[,1:10] <- lapply(df[,1:10], function(x) factor(x, levels=c("Molto in disaccordo",
                                                              "In disaccordo", "Leggermente in disaccordo", "Non saprei", "Leggermente d' accordo",
                                                              "D' accordo", "Molto d' accordo")))

df$EmpExp_Denounce <- factor(df$EmpExp_Denounce)
df$PersonalCommitment_Denounce <- factor(df$PersonalCommitment_Denounce)
}

#Missing values
{
  missmap(df)
}

#Saving the dataset in the RData format (for markdown)
save(df, file="~/Documents/Jacopo/Research_Projects/Ongoing/Tamias/MarkDown/DatasetEutamias.RData")

################################################################
#Baseline information about respondents
################################################################
{
#Location
prop.table(table(df$Location))
  
#Age
prop.table(table(df$Age, df$Location), margin = 2)
chisq.test(table(df$Age, df$Location)) #age does not differ between sites

#Sex
df$Sex <- factor(df$Sex)
prop.table(table(df$Sex))
prop.table(table(df$Sex, df$Location), margin=2)
chisq.test(table(df$Location, df$Sex)) #sex does not differ between sites

#Level of education
prop.table(table(df$Education))
prop.table(table(df$Education, df$Location), margin=2)
chisq.test(table(df$Education, df$Location)) #education varies between sites

#Frequency of visit
prop.table(table(df$Frequency_visit))
prop.table(table(df$Frequency_visit, df$Location), margin=2)
chisq.test(table(df$Frequency_visit, df$Location)) #the frequency of visits varies between sites

#Activities at the park
prop.table(table(df$Activity_birdwatching))
prop.table(table(df$Activity_birdwatching, df$Location), margin=2)

prop.table(table(df$Activity_dog))
prop.table(table(df$Activity_dog, df$Location), margin=2)

prop.table(table(df$Activity_drawingphoto))
prop.table(table(df$Activity_drawingphoto, df$Location), margin=2)

prop.table(table(df$Activity_music))
prop.table(table(df$Activity_music, df$Location), margin=2)

prop.table(table(df$Activity_picknick))
prop.table(table(df$Activity_picknick, df$Location), margin=2)

prop.table(table(df$Activity_sport))
prop.table(table(df$Activity_sport, df$Location), margin=2)

prop.table(table(df$Activity_walking))
prop.table(table(df$Activity_walking, df$Location), margin=2)

prop.table(table(df$Activity_workvolunteer))
prop.table(table(df$Activity_workvolunteer, df$Location), margin=2)

prop.table(table(df$Activity_wildifefeeding))
prop.table(table(df$Activity_wildifefeeding, df$Location), margin=2)

#Experience with squirrels
prop.table(table(df$Squirrel_watching))
prop.table(table(df$Squirrel_watching, df$Location), margin=2)
chisq.test(df$Squirrel_watching, df$Location) #No differences

prop.table(table(df$Squirrel_feeding))
prop.table(table(df$Squirrel_feeding, df$Location), margin=2)
chisq.test(df$Squirrel_feeding, df$Location) #No differences

prop.table(table(df$Squirrel_touching))
prop.table(table(df$Squirrel_touching, df$Location), margin=2)
chisq.test(df$Squirrel_touching, df$Location) #No differences

prop.table(table(df$Squirrel_exp, df$Location), margin=2)
chisq.test(table(df$Squirrel_exp, df$Location)) #H0 independence: there are no differences between areas

df$Presence_before <- factor(df$Presence_before)
prop.table(table(df$Presence_before))
prop.table(table(df$Presence_before, df$Location), margin=2)
chisq.test(table(df$Presence_before, df$Location)) #H0 independence: there are no differences between areas

#Sankey diagram for squirrel experience
nodes <- data.frame("name"= c("Rome", "Valeggio", "Heard before", "Not heard before",
                        "Observed", "Fed", "Touched"))
links <- as.data.frame(matrix(c(
  0, 2, 40, # Each row represents a link. The first number
  0, 3, 53, # represents the node being conntected from. 
  1, 2, 62, # the second number represents the node connected to.
  1, 3, 93,# The third number is the value of the node
  2, 4, 88,#
  3, 4, 35,#
  4, 5, 36,#
  5, 6, 9),#
  byrow = TRUE, ncol = 3))
names(links) <- c("source", "target", "value")
sankeyNetwork(Links = links, Nodes = nodes,
              Source = "source", Target = "target",
              Value = "value", NodeID = "name",
              fontSize= 12, nodeWidth = 30)

#How many respondents reported to have observed chipmunks, while they were unaware of
#their presence in the park
levels(df$Squirrel_watching)
idx <- c("Once", "Multiple times per month", "Multiple times per week")
nrow(df[which(df$Presence_before=="No"&df$Squirrel_watching%in%idx),])/nrow(df)

#Emotional disposition
multi.fun <- function(x) {
  c(min = mean(x), sd = sd(x))}

prop.table(table(df$Joy))
prop.table(table(df$Interest))
prop.table(table(df$Surprise))
prop.table(table(df$Fear))
prop.table(table(df$Disgust))

#Core affect
prop.table(table(df$CoreAff_spotting))

#Existence beliefs
prop.table(table(df$ExBel_future))
prop.table(table(df$ExBel_personal))

#Moral beliefs
prop.table(table(df$MoralBel))

#Empirical expectations
prop.table(table(df$EmpExp_Presence))

#Normative expectations
prop.table(table(df$NormExp_Acceptable))
names(df)

#Sanctions
prop.table(table(df$EmpExp_Denounce))
prop.table(table(df$PersonalCommitment_Denounce))
names(df)
}

#Converting likert scales to numbers
{
head(df)
str(df)
t(colnames(df))
idx <- c("Belief_envqual", "Belief_aesth", "Belief_freq", "Belief_nvisit",
         "Belief_satisf", "Belief_zoonoses", "Belief_disdogs", "Belief_veg", "Belief_nest",
         "Belief_garb", "Joy", "Fear", "Surprise", "Disgust", "Interest","ExBel_future",
         "ExBel_personal", "EmpExp_Presence", "NormExp_Acceptable") #7-points bipolar scales as numbers
df[,names(df)%in%idx] <- as.data.frame(lapply(df[,names(df)%in%idx], function(x) as.numeric(x)-4))
df$CoreAff_spotting <- as.numeric(df$CoreAff_spotting) #Unipolar scales as numbers
df$MoralBel <- as.numeric(df$MoralBel)
idx <- c("Eval_envqual", "Eval_aesth", "Eval_freq", "Eval_nvisit", "Eval_satisf", 
         "Eval_zoonoses", "Eval_disdogs", "Eval_veg", "Eval_nest", "Eval_garb")
df[,names(df)%in%idx] <- as.data.frame(lapply(df[,names(df)%in%idx], function(x)as.numeric(x)-4))
}

################################################################
#Attitudes
################################################################
{
#Reliability
{
#Overall scale (10 items per sub-dimension, unsatisfactory)
cronbach(t(do.call(rbind, df[,c(1:10)])))
cronbach(t(do.call(rbind, df[,c(39:48)])))
cronbach(t(do.call(rbind, df[,c(1:10, 39:48)])))

#McDonald's Omega reveals a disjoint structure
omega(t(do.call(rbind, df[,c(1:10)])))
omega(t(do.call(rbind, df[,c(39:48)])))
omega(t(do.call(rbind, df[,c(1:10, 39:48)])))

#And reliability of subdimension is satisfactory for the first five impacts
cronbach(t(do.call(rbind, df[,c(1:5)])))
cronbach(t(do.call(rbind, df[,c(39:43)])))
cronbach(t(do.call(rbind, df[,c(1:5, 39:43)])))
omega(t(do.call(rbind, df[,c(1:5)])))
omega(t(do.call(rbind, df[,c(39:43)])))
omega(t(do.call(rbind, df[,c(1:5, 39:43)])))

cronbach(t(do.call(rbind, df[,c(6:10)])))
cronbach(t(do.call(rbind, df[,c(44:48)])))
cronbach(t(do.call(rbind, df[,c(6:10, 44:48)])))
omega(t(do.call(rbind, df[,c(6:10)])))
omega(t(do.call(rbind, df[,c(44:48)])))
omega(t(do.call(rbind, df[,c(6:10, 44:48)])))

cronbach(t(do.call(rbind, df[,c(6,7,9,10,44,45,47,48)])))
omega(t(do.call(rbind, df[,c(6,7,9,10,44,45,47,48)])))
}

#An overall attitudinal model? (no)
{
    #Measurement model
    fullmod.attitudes <- '
    #latent variables
    belief =~ NA*Str_Aesth + Str_Freq + Str_Nvisit + Str_Zoonoses +
    Str_DisDogs + Str_Garbage
    eval=~ NA*Ev_Aesth + Ev_Freq + Ev_Nvisit + Ev_Zoonoses + 
    Ev_DisDogs + Ev_Garbage
    # residual correlations
    Str_Aesth ~~ Ev_Aesth
    Str_Freq ~~ Ev_Freq
    Str_Nvisit ~~ Ev_Nvisit
    Str_Zoonoses ~~ Ev_Zoonoses
    Str_DisDogs ~~ Ev_DisDogs
    Str_Garbage ~~ Ev_Garbage'
    
    names(df)
    idx <- c("Belief_aesth", "Belief_freq", "Belief_nvisit", "Belief_zoonoses", "Belief_disdogs", "Belief_garb",
             "Eval_aesth", "Eval_freq", "Eval_nvisit", "Eval_zoonoses", "Eval_disdogs", "Eval_garb")
    foo <- as.data.frame(lapply(df[,names(df)%in%idx], as.numeric))
    names(foo) <- c("Str_Aesth", "Str_Freq", "Str_Nvisit",
                    "Str_Zoonoses", "Str_DisDogs", "Str_Garbage",
                    "Ev_Aesth", "Ev_Freq", "Ev_Nvisit",
                    "Ev_Zoonoses", "Ev_DisDogs","Ev_Garbage")
    
    mod <- cfa(fullmod.attitudes, std.lv=TRUE, std.ov=TRUE, estimator="MLM",
               data=foo)
    
    summary(mod)
    names(fitmeasures(mod))
    mod_loads <- inspect(mod,what="std")$lambda
    mod_loads[,1]^2 #Communalities
    mod_loads[,1]^2 #Uniqueness
    mod_loads[,2]^2 #Communalities
    mod_loads[,2]^2 #Uniqueness 
    fitmeasures(mod, c("rmsea","rmsea.ci.lower", "rmsea.ci.upper",
                       "rmsea.pvalue")) #chisq is significant (the model does not fit the data well)
    fitmeasures(mod, "rmsea") #RMSEA should be lower than 0.05 (it is 0.175, it sucks)
    fitmeasures(mod, "srmr") #SRMR should be lower than 0.05 (it is 0.129, it sucks)
    fitmeasures(mod, "tli") #TLI should be greater than 0.9 (it is 0.608, it sucks)
    fitmeasures(mod, "cfi") #CFI should be higher than 0.95 (it is 0.721, not so well)
    fitmeasures(mod, c("chisq", "df", "pvalue"))
}

#First five beliefs (Environmental quality, aesthetics, frequency, n.visitors, satisfaction)
#McDonald's omega raises some doubts about satisfaction and environmental quality
{
#Full model
#Measurement model
fullmod.FirstFive <- '
  #latent variables
  belief =~ NA*Str_EnvQual + Str_Aesth + Str_Freq + Str_Nvisit + Str_Satisf
  eval=~ NA*Ev_EnvQual + Ev_Aesth + Ev_Freq + Ev_Nvisit + Ev_Satisf
  # residual correlations
    Str_EnvQual ~~ Ev_EnvQual
    Str_Aesth ~~ Ev_Aesth
    Str_Freq ~~ Ev_Freq
    Str_Nvisit ~~ Ev_Nvisit
    Str_Satisf ~~ Ev_Satisf'

#Preparing the dataset
foo <- as.data.frame(lapply(df[,c(1:5, 39:43)], as.numeric))
names(foo) <- c("Str_EnvQual", "Str_Aesth", "Str_Freq", "Str_Nvisit", "Str_Satisf",
                "Ev_EnvQual", "Ev_Aesth", "Ev_Freq", "Ev_Nvisit", "Ev_Satisf")

#Estimator for non-normal continous variables (good for Likert scales, outperforms WMLS estimator)

m1.FirstFive <- cfa(fullmod.FirstFive, std.lv=TRUE, std.ov=TRUE, estimator="MLM",
                    data=foo)

summary(m1.FirstFive)
fitmeasures(m1.FirstFive)
m1.FirstFive_loads <- inspect(m1.FirstFive,what="std")$lambda
m1.FirstFive_loads[,1]^2 #Communalities
m1.FirstFive_loads[,1]^2 #Uniqueness
m1.FirstFive_loads[,2]^2 #Communalities
1-m1.FirstFive_loads[,2]^2 #Uniqueness 
fitmeasures(m1.FirstFive, c("chisq", "df", "pvalue")) #chisq is significant (the model does not fit the data well)
fitmeasures(m1.FirstFive, c("rmsea","rmsea.ci.lower", "rmsea.ci.upper",
            "rmsea.pvalue")) #RMSEA should be lower than 0.09 (it is 0.099 and p<0.05, so it does not fit well)
fitmeasures(m1.FirstFive, "srmr") #SRMR should be lower than 0.05 (it is 0.042)
fitmeasures(m1.FirstFive, "tli") #TLI should be greater than 0.9 (it is 0.948)
fitmeasures(m1.FirstFive, "cfi") #CFI should be higher than 0.95 (it is 0.966)

#No environmental quality
omega(t(do.call(rbind, lapply((df[which(!is.na(df[,c(2:5,40:43)])),c(2:5,40:43)]), as.numeric))))

nested1.FirstFive <- '
  #latent variables
belief =~ NA*Str_Aesth + Str_Freq + Str_Nvisit + Str_Satisf
eval=~ NA*Ev_Aesth + Ev_Freq + Ev_Nvisit + Ev_Satisf
# residual correlations
Str_Aesth ~~ Ev_Aesth
Str_Freq ~~ Ev_Freq
Str_Nvisit ~~ Ev_Nvisit
Str_Satisf ~~ Ev_Satisf'

#Preparing the dataset
foo <- as.data.frame(lapply(df[,c(2:5, 40:43)], as.numeric))
names(foo) <- c("Str_Aesth", "Str_Freq", "Str_Nvisit", "Str_Satisf",
                "Ev_Aesth", "Ev_Freq", "Ev_Nvisit", "Ev_Satisf")

m2.FirstFive <- cfa(nested1.FirstFive, std.lv=TRUE, std.ov=TRUE, estimator="MLM",
                    data=foo)

summary(m2.FirstFive)
fitmeasures(m2.FirstFive)
m2.FirstFive_loads <- inspect(m2.FirstFive,what="std")$lambda
m2.FirstFive_loads[,1]^2 #Communalities
m2.FirstFive_loads[,1]^2 #Uniqueness
m2.FirstFive_loads[,2]^2 #Communalities
1-m2.FirstFive_loads[,2]^2 #Uniqueness 
fitmeasures(m2.FirstFive, c("chisq", "df", "pvalue"), estimator="MLM") #chisq is significant (the model does not fit the data well)
fitmeasures(m2.FirstFive, c("rmsea","rmsea.ci.lower", "rmsea.ci.upper",
            "rmsea.pvalue")) #RMSEA should be lower than 0.05 (it is 0.099 and p<0.05, so it does not fit well)
fitmeasures(m2.FirstFive, "srmr") #SRMR should be lower than 0.05 (it is 0.039)
fitmeasures(m2.FirstFive, "tli") #TLI should be greater than 0.9 (it is 0.95)
fitmeasures(m2.FirstFive, "cfi") #CFI should be higher than 0.95 (it is 0.973)

#Dropping satisfaction
nested2.FirstFive <- '
  #latent variables
belief =~ NA*Str_Aesth + Str_Freq + Str_Nvisit
eval=~ NA*Ev_Aesth + Ev_Freq + Ev_Nvisit
# residual correlations
Str_Aesth ~~ Ev_Aesth
Str_Freq ~~ Ev_Freq
Str_Nvisit ~~ Ev_Nvisit'

#Preparing the dataset
names(df)
foo <- as.data.frame(lapply(df[,c(2:4, 40:42)], as.numeric))
names(foo) <- c("Str_Aesth", "Str_Freq", "Str_Nvisit",
                "Ev_Aesth", "Ev_Freq", "Ev_Nvisit")

m3.FirstFive <- cfa(nested2.FirstFive, std.lv=TRUE, std.ov=TRUE, estimator="MLM",
                    data=foo)

summary(m3.FirstFive)
fitmeasures(m3.FirstFive)
m3.FirstFive_loads <- inspect(m3.FirstFive,what="std")$lambda
m3.FirstFive_loads[,1]^2 #Communalities
m3.FirstFive_loads[,2]^2 #Communalities
fitmeasures(m3.FirstFive, c("chisq", "df", "pvalue"), estimator="MLM") #chisq is significant (the model does not fit the data well)
fitmeasures(m3.FirstFive, c("rmsea","rmsea.ci.lower", "rmsea.ci.upper",
            "rmsea.pvalue")) #RMSEA should be lower than 0.05 (it is 0.075 and p>0.05, so the model is nice)
fitmeasures(m3.FirstFive, "srmr") #SRMR should be lower than 0.05 (it is 0.035)
fitmeasures(m3.FirstFive, "tli") #TLI should be greater than 0.9 (it is 0.97)
fitmeasures(m3.FirstFive, "cfi") #CFI should be higher than 0.95 (it is 0.991)

#Likelihood ration testing and model comparisons
c(fitmeasures(m1.FirstFive, c("srmr")) ,fitmeasures(m2.FirstFive, c("srmr")), 
  fitmeasures(m3.FirstFive, c("srmr"))) #SRMR should be lower than 0.05
c(fitmeasures(m1.FirstFive, c("rmr")) ,fitmeasures(m2.FirstFive, c("rmr")),
  fitmeasures(m3.FirstFive, c("rmr"))) #RootMean Square Residual, the lower the better
c(fitmeasures(m1.FirstFive, c("tli")) ,fitmeasures(m2.FirstFive, c("tli")),
  fitmeasures(m3.FirstFive, c("tli"))) #TLI should be greater than 0.9
c(fitmeasures(m1.FirstFive, c("cfi")) ,fitmeasures(m2.FirstFive, c("cfi")),
  fitmeasures(m3.FirstFive, c("cfi"))) #CFI should be higher than 0.95
c(fitmeasures(m1.FirstFive, c("ifi")) ,fitmeasures(m2.FirstFive, c("ifi")),
  fitmeasures(m3.FirstFive, c("ifi"))) #IFI Value closer to one indicate better fit
c(fitmeasures(m1.FirstFive, c("ecvi")) ,fitmeasures(m2.FirstFive, c("ecvi")),
  fitmeasures(m3.FirstFive, c("ecvi"))) #Expected cross-validation index
c(fitmeasures(m1.FirstFive, c("bic")) ,fitmeasures(m2.FirstFive, c("bic")),
  fitmeasures(m3.FirstFive, c("bic"))) #Bayesian Information Criteria
c(fitmeasures(m1.FirstFive, c("aic")) ,fitmeasures(m2.FirstFive, c("aic")),
  fitmeasures(m3.FirstFive, c("aic"))) #Akaike's Information Criteria
c(fitmeasures(m1.FirstFive, c("bic2")) ,fitmeasures(m2.FirstFive, c("bic2")),
  fitmeasures(m3.FirstFive, c("bic2"))) #Sample-adjusted BIC
c(fitmeasures(m1.FirstFive, c("gfi")) ,fitmeasures(m2.FirstFive, c("gfi")),
  fitmeasures(m3.FirstFive, c("gfi"))) #GoF index
c(fitmeasures(m1.FirstFive, c("agfi")) ,fitmeasures(m2.FirstFive, c("agfi")),
  fitmeasures(m3.FirstFive, c("agfi"))) #Adjusted GoF index
c(fitmeasures(m1.FirstFive, c("pgfi")) ,fitmeasures(m2.FirstFive, c("pgfi")),
  fitmeasures(m3.FirstFive, c("pgfi"))) #Adjusted GoF index
c(fitmeasures(m1.FirstFive, c("mfi")) ,fitmeasures(m2.FirstFive, c("mfi")),
  fitmeasures(m3.FirstFive, c("mfi"))) #Adjusted GoF index
anova(m1.FirstFive, m2.FirstFive, m3.FirstFive) #Dropping environmental quality improves the model

#Residual: "a careful examination of the residuals may reveal an unsystematic scatter across the matrix
#with no troublingly large values. Clusters of residuals, which suggest and could support additional
#factors defined by three or more measurest. Indicatively, a residual less than .1 would not allow a
#product of two salient loadings (>.3)
residuals(m3.FirstFive)$cov #Fine

#Summarizing model output (LV scores, variances, covariances)
summary(m3.FirstFive)

#Plotting the best model
semPaths(m3.FirstFive, intercepts=FALSE, title = FALSE, layout = "tree2")

#Fig. 3
semPaths(m1.FirstFive, intercepts=FALSE, title = FALSE, layout = "tree2")

}

#Last five beliefs (Zoonoses, disease to dogs, nest predation, vegetation damaging, garbage)
#McDonald's omega raises some doubts about vegetation damaging and nest predation
{
#Measurement model
fullmod.LastFive <- '
  #latent variables
  belief =~ NA*Str_Zoonoses + Str_DisDogs + Str_VegDam + Str_NestPred + Str_Garbage
  eval=~ NA*Ev_Zoonoses + Ev_DisDogs + Ev_VegDam + Ev_NestPred + Ev_Garbage
  # residual correlations
    Str_Zoonoses ~~ Ev_Zoonoses
    Str_DisDogs ~~ Ev_DisDogs
    Str_VegDam ~~ Ev_VegDam
    Str_NestPred ~~ Ev_NestPred
    Str_Garbage ~~ Ev_Garbage'

#Estimator for non-normal continous variables (good for Likert scales)
foo <- as.data.frame(lapply(df[,c(6:10, 44:48)], as.numeric))
names(foo) <- c("Str_Zoonoses", "Str_DisDogs", "Str_VegDam", "Str_NestPred", "Str_Garbage",
                "Ev_Zoonoses", "Ev_DisDogs", "Ev_VegDam", "Ev_NestPred", "Ev_Garbage")

m1.LastFive <- cfa(fullmod.LastFive, std.lv=TRUE, std.ov=TRUE, estimator="MLM",
                        data=foo)

summary(m1.FirstFive)
fitmeasures(m1.FirstFive)
m1.LastFive_loads <- inspect(m1.LastFive,what="std")$lambda
m1.LastFive_loads[,1]^2 #Communalities
m1.LastFive_loads[,1]^2 #Uniqueness
m1.LasttFive_loads[,2]^2 #Communalities
1-m1.LastFive_loads[,2]^2 #Uniqueness 
fitmeasures(m1.LastFive, c("rmsea","rmsea.ci.lower", "rmsea.ci.upper",
                           "rmsea.pvalue")) #chisq is significant (the model does not fit the data well)
fitmeasures(m1.LastFive, "rmsea") #RMSEA should be lower than 0.05 (it is 0.08, not so well)
fitmeasures(m1.LastFive, "srmr") #SRMR should be lower than 0.05 (it is 0.06, not so well)
fitmeasures(m1.LastFive, "tli") #TLI should be greater than 0.9 (it is 0.88, not so well)
fitmeasures(m1.LastFive, "cfi") #CFI should be higher than 0.95 (it is 0.92, not so well)

#The fis is unsatisfactory, however omega reveals that impacts on vegetation and nest predation might compromise the scale
#let's try to remove them and re-fit the scale.

nested1.LastFive <- '
  #latent variables
belief =~ NA*Str_Zoonoses + Str_DisDogs + Str_NestPred + Str_Garbage
eval=~ NA*Ev_Zoonoses + Ev_DisDogs + Ev_NestPred + Ev_Garbage
# residual correlations
Str_Zoonoses ~~ Ev_Zoonoses
Str_DisDogs ~~ Ev_DisDogs
Str_NestPred ~~ Ev_NestPred
Str_Garbage ~~ Ev_Garbage'

#Estimator for non-normal continous variables (good for Likert scales)
foo <- as.data.frame(lapply(df[,c(6,7,9,10, 44,45,47,48)], as.numeric))
names(foo) <- c("Str_Zoonoses", "Str_DisDogs", "Str_NestPred", "Str_Garbage",
                "Ev_Zoonoses", "Ev_DisDogs", "Ev_NestPred", "Ev_Garbage")

m2.LastFive <- cfa(nested1.LastFive, std.lv=TRUE, std.ov=TRUE, estimator="MLM",
                        data=foo)

summary(m2.LastFive)
fitmeasures(m2.LastFive)
m2.LastFive_loads <- inspect(m2.LastFive,what="std")$lambda
m2.LastFive_loads[,1]^2 #Communalities
m2.LastFive_loads[,1]^2 #Uniqueness
m2.LastFive_loads[,2]^2 #Communalities
1-m2.LastFive_loads[,2]^2 #Uniqueness 
fitmeasures(m2.LastFive, c("rmsea","rmsea.ci.lower", "rmsea.ci.upper",
                            "rmsea.pvalue")) #chisq is non significant (the model is not so bad)
fitmeasures(m2.LastFive, "srmr") #SRMR should be lower than 0.05 (it is 0.055, borderline)
fitmeasures(m2.LastFive, "tli") #TLI should be greater than 0.9 (it is 0.936, nice)
fitmeasures(m2.LastFive, "cfi") #CFI should be higher than 0.95 (it is 0.966, nice)

#Dropping nest damaging

nested1.LastFive <- '
  #latent variables
belief =~ NA*Str_Zoonoses + Str_DisDogs + Str_Garbage
eval=~ NA*Ev_Zoonoses + Ev_DisDogs + Ev_Garbage
# residual correlations
Str_Zoonoses ~~ Ev_Zoonoses
Str_DisDogs ~~ Ev_DisDogs
Str_Garbage ~~ Ev_Garbage'

#Estimator for non-normal continous variables (good for Likert scales)
foo <- as.data.frame(lapply(df[,c(6,7,10, 44,45,48)], as.numeric))
names(foo) <- c("Str_Zoonoses", "Str_DisDogs", "Str_Garbage",
                "Ev_Zoonoses", "Ev_DisDogs", "Ev_Garbage")

m3.LastFive <- cfa(nested1.LastFive, std.lv=TRUE, std.ov=TRUE, estimator="MLM",
                        data=foo)

summary(m3.LastFive)
fitmeasures(m3.LastFive)
m3.LastFive_loads <- inspect(m3.LastFive,what="std")$lambda
m3.LastFive_loads[,1]^2 #Communalities
m3.LastFive_loads[,1]^2 #Uniqueness
m3.LastFive_loads[,2]^2 #Communalities
1-m3.LastFive_loads[,2]^2 #Uniqueness 
fitmeasures(m3.LastFive, c("rmsea","rmsea.ci.lower", "rmsea.ci.upper",
                            "rmsea.pvalue")) #chisq is non significant (the model is not so bad)
fitmeasures(m3.LastFive, "srmr") #SRMR should be lower than 0.05 (it is 0.035, nice)
fitmeasures(m3.LastFive, "tli") #TLI should be greater than 0.9 (it is 0.974, nice)
fitmeasures(m3.LastFive, "cfi") #CFI should be higher than 0.9 (it is 0.991, nice)

#Likelihood ration testing and model comparisons
c(fitmeasures(m1.LastFive, c("srmr")) ,fitmeasures(m2.LastFive, c("srmr")),
  fitmeasures(m3.LastFive, c("srmr"))) #SRMR should be lower than 0.05
c(fitmeasures(m1.LastFive, c("rmr")) ,fitmeasures(m2.LastFive, c("rmr")),
  fitmeasures(m3.LastFive, c("rmr"))) #RootMean Square Residual, the lower the better
c(fitmeasures(m1.LastFive, c("tli")) ,fitmeasures(m2.FirstFive, c("tli")),
  fitmeasures(m3.LastFive, c("tli"))) #TLI should be greater than 0.9
c(fitmeasures(m1.LastFive, c("cfi")) ,fitmeasures(m2.FirstFive, c("cfi")),
  fitmeasures(m3.LastFive, c("cfi"))) #CFI should be higher than 0.95
c(fitmeasures(m1.LastFive, c("ifi")) ,fitmeasures(m2.FirstFive, c("ifi")),
  fitmeasures(m3.LastFive, c("ifi"))) #IFI Value closer to one indicate better fit
c(fitmeasures(m1.LastFive, c("ecvi")) ,fitmeasures(m2.FirstFive, c("ecvi")),
  fitmeasures(m3.LastFive, c("ecvi"))) #Expected cross-validation index
c(fitmeasures(m1.LastFive, c("bic")) ,fitmeasures(m2.FirstFive, c("bic")),
  fitmeasures(m3.LastFive, c("bic"))) #Bayesian Information Criteria
c(fitmeasures(m1.LastFive, c("aic")) ,fitmeasures(m2.FirstFive, c("aic")),
  fitmeasures(m3.LastFive, c("aic"))) #Akaike's Information Criteria
c(fitmeasures(m1.LastFive, c("bic2")) ,fitmeasures(m2.FirstFive, c("bic2")),
  fitmeasures(m3.LastFive, c("bic2"))) #Sample-adjusted BIC
c(fitmeasures(m1.LastFive, c("gfi")) ,fitmeasures(m2.FirstFive, c("gfi")),
  fitmeasures(m3.LastFive, c("gfi"))) #GoF index
c(fitmeasures(m1.LastFive, c("agfi")) ,fitmeasures(m2.FirstFive, c("agfi")),
  fitmeasures(m3.LastFive, c("agfi"))) #Adjusted GoF index
c(fitmeasures(m1.LastFive, c("pgfi")) ,fitmeasures(m2.FirstFive, c("pgfi")),
  fitmeasures(m3.LastFive, c("pgfi"))) #Adjusted GoF index
c(fitmeasures(m1.LastFive, c("mfi")) ,fitmeasures(m2.FirstFive, c("mfi")),
  fitmeasures(m3.LastFive, c("mfi"))) #Adjusted GoF index
anova(m3.LastFive, m2.LastFive, m1.LastFive) #Dropping nest damages slightly improves the model

#Plotting residuals
residuals(m3.LastFive)$cov #Garbage and dog diseases might be correlated

#Summarizing model output (LV scores, variances, covariances)
summary(m3.LastFive)

#Plotting best model
semPaths(m3.LastFive, intercepts=FALSE, title = FALSE, layout = "tree2")

}

#B*e scoring
{
names(df)
idx <- c("Belief_aesth", "Belief_freq", "Belief_nvisit", "Belief_zoonoses", "Belief_disdogs", "Belief_garb")
idx.2 <- c("Eval_aesth", "Eval_freq", "Eval_nvisit", "Eval_zoonoses", "Eval_disdogs", "Eval_garb")
foo <- as.data.frame(lapply(df[,names(df)%in%idx], function(x) as.numeric(x)-4))
fee <- as.data.frame(lapply(df[,names(df)%in%idx.2], function(x) as.numeric(x)))
faa <- data.frame((foo$Belief_aesth*fee$Eval_aesth), (foo$Belief_freq*fee$Eval_freq),
                  (foo$Belief_nvisit*fee$Eval_nvisit))
names(faa) <- c("Aesth", "Freq", "Nvisit")
att.score <- apply(faa,1,sum)
df$Score_attPositiveImpacts <- att.score
hist(df$Score_attPositiveImpacts)
boxplot(Score_attPositiveImpacts ~ Location, data=df)

names(df)
faa <- data.frame((foo$Belief_zoonoses*fee$Eval_zoonoses), (foo$Belief_disdogs*fee$Eval_disdogs),
                  (foo$Belief_garb*fee$Eval_garb))
names(faa) <- c("Zoonoses", "DisDogs", "Garbage")
att.score <- apply(faa,1,sum)
df$Score_attNegativeImpacts <- att.score

par(mfrow=c(1,2))
hist(df$Score_attNegativeImpacts)
hist(df$Score_attPositiveImpacts)
par(mfrow=c(1,1))}
}

################################################################
#Is it possible to segment respondents?
################################################################
{
#comparing K-means, hierarchical and PAM (K-medoids) cluster analysis
idx <- c("Joy", "Fear", "Surprise", "Disgust", "Interest", "ExBel_future", "ExBel_personal",
         "MoralBel", "EmpExp_Presence", "NormExp_Acceptable","Score_attNegativeImpacts", 
         "Score_attPositiveImpacts", "CoreAff_spotting")
dat <- df[,names(df)%in%idx]
dat$ID <- rownames(dat)
missing.idx <- dat[!complete.cases(dat), "ID"]
cluster.dat <- dat[complete.cases(dat),]
cluster.dat <- cluster.dat[,-14] #Removing IDs
cluster.dat <- as.data.frame(lapply(cluster.dat, function(x) scale(x, center=TRUE, scale=TRUE)))

#Clustering tendency: Hopkins index
head(cluster.dat)
set.seed(123)
hopkins(cluster.dat, n = nrow(cluster.dat)-1)

random_df <- apply(cluster.dat, 2,
                   function(x){runif(length(x), min(x), (max(x)))})
random_df <- as.data.frame(random_df)
set.seed(123)
hopkins(random_df, n = nrow(random_df)-1)

#Comparing clustering algorithms
clmethods <- c("hierarchical","kmeans","pam")
intern <- clValid(cluster.dat, nClust = 2:8,
                  clMethods = clmethods, validation = "internal")
stab <- clValid(cluster.dat, nClust = 2:8, clMethods = clmethods,
                validation = "stability")
summary(intern)
optimalScores(intern)
optimalScores(stab)

#No method outperforms the other ones. This might indicate the lack of any real cluster
#Let's try to estimate p-values in a hierarchical cluster analysis
set.seed(123)
pval.hclust <- pvclust(cluster.dat, method.dist="euclidean",
                  method.hclust="complete", nboot = 10)
plot(pval.hclust, hang = -1, cex = 0.5)
pvrect(pval.hclust)
pval.hclust
}

################################################################
#Comparing areas
################################################################

#Do attitudes score differ between different the two areas and between groups of users?
{
  df$Location <- factor(df$Location)
  df$Squirrel_exp <- factor(df$Squirrel_exp, labels=c("No", "Yes"))
  
  hist(df$Score_attNegativeImpacts, breaks=20)
  mod_AttNeg <- glm(Score_attNegativeImpacts ~ Location * Squirrel_exp, data=df, family=gaussian)
  summary(mod_AttNeg)
  plot_model(mod_AttNeg, type="eff", terms="Location")
  plot_model(mod_AttNeg, type="eff", terms="Squirrel_exp")
  plot_model(mod_AttNeg, type="int", title="Negative impacts (garbage, zoonoses, disease to dogs)", 
             show.legend=FALSE)
  
  hist(df$Score_attPositiveImpacts, breaks=20)
  mod_AttPos <- glm(Score_attPositiveImpacts ~ Location * Squirrel_exp, data=df, family=gaussian)
  summary(mod_AttPos)
  plot_model(mod_AttPos, type="eff", terms="Location")
  plot_model(mod_AttPos, type="eff", terms="Squirrel_exp")
  plot_model(mod_AttPos, type="int", title="Negative impacts (garbage, zoonoses, disease to dogs)", 
             show.legend=FALSE)
  
}

#Does core affect differ between different the two areas and between groups of users?
{
  barplot(table(df$CoreAff_spotting))
  mod_CoreAff <- glm(CoreAff_spotting ~ Location * Squirrel_exp, data=df, family=Gamma(link="identity"))
  summary(mod_CoreAff)
  plot_model(mod_CoreAff, type="eff", terms="Location")
  plot_model(mod_CoreAff, type="eff", terms="Squirrel_exp")
  plot_model(mod_CoreAff, type="int",
             title="Core affect", show.legend=FALSE)
}

#Does existence beliefs vary between areas and groups of respondents?
{
  hist(df$ExBel_future)
  foo <- df
  foo$ExBel_future <-   foo$ExBel_future + 4 
  modExBel_future <- glm(ExBel_future ~ Location * Squirrel_exp, data=foo, family=Gamma(link="identity"))
  summary(modExBel_future)
  plot_model(modExBel_future, type="eff", terms="Location")
  plot_model(modExBel_future, type="eff", terms="Squirrel_exp")
  plot_model(modExBel_future, type="int", 
             title="Importance for the future generations", show.legend=FALSE)
  
  foo$ExBel_personal <-   foo$ExBel_personal + 4 
  hist(df$ExBel_personal)
  modExBel_personal <- glm(as.numeric(ExBel_personal) ~ Location * Squirrel_exp, data=foo,
                           family=Gamma(link="identity"))
  summary(modExBel_personal)
  plot_model(modExBel_personal, type="eff", terms="Location")
  plot_model(modExBel_personal, type="eff", terms="Squirrel_exp")
  plot_model(modExBel_personal, type="int")
}

#Does normative beliefs differ between different the two areas and between groups of users?
{
  modMoralBel <- glm(as.numeric(MoralBel) ~ Location * Squirrel_exp, data=df, family=Gamma(link="identity"))
  summary(modMoralBel)
  plot_model(modMoralBel, type="eff", terms="Location")
  plot_model(modMoralBel, type="eff", terms="Squirrel_exp")
  plot_model(modMoralBel, type="int")
  
  foo$EmpExp_Presence <- foo$EmpExp_Presence + 4
  modEmpExp <- glm(as.numeric(EmpExp_Presence) ~ Location * Squirrel_exp, data=foo, family=Gamma(link="identity"))
  summary(modEmpExp)
  plot_model(modEmpExp, type="eff", terms="Location")
  plot_model(modEmpExp, type="eff", terms="Squirrel_exp")
  plot_model(modEmpExp, type="int")
  
  #Does normative expectations
  df <- plyr::rename(df, c("NormExp_Acceptable" = "Normative_expectations"))
  df$Normative_expectations <- df$Normative_expectations + 4
  modNormExp <- glm(Normative_expectations ~ Location * Squirrel_exp, data=df, family=Gamma(link="identity"))
  summary(modNormExp)
  plot_model(modNormExp, type="eff", terms="Location")
  plot_model(modNormExp, type="eff", terms="Squirrel_exp")
  plot_model(modNormExp, type="int", show.legend=FALSE)
}

#Multiple plots (Fig.4)
{
  require(gridExtra)
  grid.arrange(pA, pB, nrow = 1)
}

################################################################
#Geographic profiling
################################################################
require(RgeoProfile)
require(stringr)
require(ggmap)
register_google(key ="AIzaSyDV9QP7CdmQ3YApschivng59DoRY3rs8Rg")
require(foreign)

{
#Roma
{
roma.df <- read.csv(file="~/Documents/Jacopo/Research_Projects/Ongoing/Tamias/Data/GeoProfile/Villa-Ada-coordinate_UTM-Totale.csv")
head(roma.df)

roma.df$lat <- str_replace(roma.df$lat, ",", ".")
roma.df$lat <- as.numeric(roma.df$lat)
roma.df$long <- str_replace(roma.df$long, ",", ".")
roma.df$long <- as.numeric(roma.df$long)

disp.dist <- 1 #Dispersal parameter
d_Roma <- geoData(roma.df$long, roma.df$lat)
p_Roma <- geoParams(data = d_Roma, sigma_mean = disp.dist, sigma_squared_shape = 2, chains = 3, 
               burnin = 1e3, samples = 1e4, longitude_minMax = c(12.490, 12.513167),
               latitude_minMax = c(41.9218, 41.939))
m_Roma <- geoMCMC(data = d_Roma, params = p_Roma, lambda=0.05) #run the MCMC model
gp_Roma <- geoProfile(m_Roma$posteriorSurface)
p1 <- geoPlotSigma(params = p_Roma, mcmc = m_Roma) # plot prior and posterior of sigma
p1

#Probabilistic profiling
mapRoma <- geoPlotMap(params = p_Roma, data = d_Roma, surface = m_Roma$geoProfile,
           breakPercent = seq(0, 50, 5), gpLegend=FALSE, mapType = "hybrid",
           crimeCol = "black", crimeBorderCol="white", crimeBorderWidth = 1.5, crimeCex = 3, sourceCol = "red", sourceCex = 2, 
           latLimits= c(41.9218, 41.939), lonLimits=c(12.490, 12.513167))# profile on the map
mapRoma

#Plot contours, crime and sources without map
contour(t(m_Roma$geoProfile), x=seq(min(d_Roma$longitude), max(d_Roma$longitude), 
        length.out=nrow(m_Roma$geoProfile)),
        y=seq(min(d_Roma$latitude), max(d_Roma$latitude), length.out=ncol(m_Roma$geoProfile)))
points(cbind(d_Roma$longitude, d_Roma$latitude), col = "red", pch = 16)

}

#Valeggio
{
valeggio.df <- read.dbf(file="~/Documents/Jacopo/Research_Projects/Ongoing/Spatial_Profiling/Data/Tamias/Shapefile/Tamias_Valleggio.dbf")
foo <- valeggio.df
sort(foo$lat)
table(foo$lat)
which(foo$lat>45.36570)
#foo[22,"lat"] <- 45.36209
which(foo$lat<45.35062)
#foo[68,"lat"] <- 45.35896
sort(foo$lat)
valeggio.df <- foo
range(valeggio.df$lat)

#Dividing into 5-years period
valeggio.df$period <- NULL
valeggio.df[which(valeggio.df$year_first<=2002), "period"] <- "1997-2002"
valeggio.df[which(valeggio.df$year_first%in%2003:2008), "period"] <- "2003-2008"
valeggio.df[which(valeggio.df$year_first%in%2009:2013), "period"] <- "2009-2013"
valeggio.df[which(valeggio.df$year_first%in%2014:2018), "period"] <- "2014-2018"

#Setting up dispersal distance (max 200 m in Tamias sibiricus)
disp.dist <- 1 #Dispersal parameter

d_Valeggio <- geoData(valeggio.df$long, valeggio.df$lat)
p_Valeggio <- geoParams(data = d_Valeggio, sigma_mean = disp.dist, sigma_squared_shape = 2, chains = 3, 
               burnin = 1e3, samples = 1e4, longitude_minMax = c(10.718, 10.743),
               latitude_minMax = c(45.348, 45.3655))
m_Valeggio <- geoMCMC(data = d_Valeggio, params = p_Valeggio, lambda=0.05) #run the MCMC model
gp <- geoProfile(m_Valeggio$posteriorSurface)
p2 <- geoPlotSigma(params = p_Valeggio, mcmc = m_Valeggio) # plot prior and posterior of sigma


#Probabilistic profile
mapValeggio <- geoPlotMap(params = p_Valeggio, data = d_Valeggio, surface = m_Valeggio$geoProfile,
              breakPercent = seq(0, 50, 5), gpLegend=FALSE, mapType = "hybrid",
              crimeCol = "black", crimeBorderCol="white", crimeBorderWidth = 1.5, crimeCex = 3, 
              sourceCol = "red", sourceCex = 2, plotContours = TRUE, smoothScale = FALSE,
              latLimits= c(45.348, 45.3655), lonLimits=c(10.718, 10.743), zoom=NULL)# profile on the map
mapValeggio

#plot contours, crime and sources without map
contour(t(m_Valeggio$geoProfile), x=seq(min(d_Valeggio$longitude), max(d_Valeggio$longitude),
                                        length.out=nrow(m_Valeggio$geoProfile)), y=seq(min(d_Valeggio$latitude), max(d_Valeggio$latitude),
                                                                                       length.out=ncol(m_Valeggio$geoProfile)))
points(cbind(d_Valeggio$longitude, d_Valeggio$latitude), col = "red", pch = 16)

}

#Ad-hoc function
{
line2user <- function(line, side) {
  lh <- par('cin')[2] * par('cex') * par('lheight')
  x_off <- diff(grconvertX(0:1, 'inches', 'user'))
  y_off <- diff(grconvertY(0:1, 'inches', 'user'))
  switch(side,
         `1` = par('usr')[3] - line * y_off * lh,
         `2` = par('usr')[1] - line * x_off * lh,
         `3` = par('usr')[4] + line * y_off * lh,
         `4` = par('usr')[2] + line * x_off * lh,
         stop("side must be 1, 2, 3, or 4", call.=FALSE))
}
}

#Creating a joint plot of dispersal and source-sink dynamics
{
par(mfrow=c(2,2))
geoPlotSigma(params = p_Roma, mcmc = m_Roma) # plot prior and posterior of sigma
contour(t(m_Roma$geoProfile), x=seq(min(d_Roma$longitude), max(d_Roma$longitude), 
        length.out=nrow(m_Roma$geoProfile)),
        y=seq(min(d_Roma$latitude), max(d_Roma$latitude), length.out=ncol(m_Roma$geoProfile)))
points(cbind(d_Roma$longitude, d_Roma$latitude), col = "red", pch = 16)
text(line2user(line=mean(par('mar')[c(2, 4)]), side=2), 
     line2user(line=2, side=3), 'Rome', xpd=NA, cex=1.5, font=2)
geoPlotSigma(params = p_Valeggio, mcmc = m_Valeggio) # plot prior and posterior of sigma
contour(t(m_Valeggio$geoProfile), x=seq(min(d_Valeggio$longitude), max(d_Valeggio$longitude),
      length.out=nrow(m_Valeggio$geoProfile)), y=seq(min(d_Valeggio$latitude), max(d_Valeggio$latitude),
      length.out=ncol(m_Valeggio$geoProfile)))
points(cbind(d_Valeggio$longitude, d_Valeggio$latitude), col = "red", pch = 16)
text(line2user(line=mean(par('mar')[c(2, 4)]), side=2), 
     line2user(line=2, side=3), 'Valeggio sul Mincio', xpd=NA, cex=1.5, font=2)
par(mfrow=c(1,1))
}
}
```
